# On the relationship between concentration buffering and noise reduction in phase separating systems

**DOI:** 10.1101/2024.02.26.579621

**Authors:** Christoph Zechner, Frank Jülicher

## Abstract

Biomolecular condensates have been proposed to buffer intracellular concentrations and reduce noise. Recent results demonstrate that concentrations need not be buffered in multicomponent systems, leading to a non-constant saturation concentration ( *c*_*sat*_ ) when individual components are varied. Simplified equilibrium considerations suggest that noise reduction might be closely related to concentration buffering and that a fixed saturation concentration is required for noise reduction to be effective. Here we present a theoretical analysis to demonstrate that these suggestions do not apply to mesoscopic fluctuating systems. We show that concentration buffering and noise reduction are distinct concepts, which cannot be used interchangeably. We further demonstrate that concentration buffering – and a constant *c*_*sat*_ – are neither necessary nor sufficient for noise reduction to be effective. Clarity about these concepts is important for studying the role of condensates in controlling cellular noise and for the interpretation of concentration relationships in cells.

## Introduction

Biomolecular condensates are membraneless compartments that are segregated from the surrounding cytoplasm by phase coexistence^1^. The thermodynamic constraints of coexisting phases restrict the potential concentration ranges of multicomponent mixtures^2-8^. This opens the possibility for cells to regulate concentrations by condensate formation. An important question is how such physical processes can be utilized to control fluctuations and molecular noise^1,9,10^.

We have previously demonstrated noise reduction by condensates theoretically as well as experimentally using engineered and endogenous systems^11^. Others have observed that the coexisting concentrations of certain intracellular condensates are not buffered when individual components are overexpressed^7^. In other words, the saturation concentrations are variable^7^. Some studies have considered concentration buffering and noise reduction as interchangable^7,12,13^, suggesting that a variable saturation concentration hampers noise reduction. However, noise reduction has been demonstrated for an endogenously labelled component of the nucleolus^11^, an intracellular condensate exhibiting a variable saturation concentration^7^. This raises the question in what situations do noise reduction and concentration buffering become related and when these phenomena are clearly distinct. Here we provide precise definitions of the concepts of concentration buffering and noise reduction in and out of equilibrium and present theory that clarifies if and how they relate to each other. To illustrate these concepts, we will discuss them in the context of previously published data from synthetic and endogenous condensates in cells^7,11^.

### Concentration buffering

We consider a multicomponent system with two coexisting phases that differ in the concentrations of their components (**Fig. 1a**). We focus on one component *A* that partitions between the two phases and denote by *c* the total average concentration *c* = a/*V*, i.e., the total copy number of molecules *a* per volume *V* of both phases combined. The concentrations of the same component in the coexisting phases are referred to as *c*_1_ and *c*_2_, respectively. We refer to *c*_1_ and *c*_2_ as dilute- and dense phase concentration, where *c*_2_ > c_1_. The partitioning of the component *A* is generally composition-dependent and therefore, *c*_1_ and *c*_2_ depend on *c* and on the total concentrations of all other components. In the following, we focus on the relationship between dilute phase concentration *c*_1_ and total concentration *c* and consider the total concentrations of all other components to be fixed. When the system is in thermodynamic equilibrium and taking the thermodynamic limit, the dilute phase concentration *c*_1_ is described by the binodal manifold in a multicomponent phase diagram. It can be formally expressed as a unique function *c*_1_ = *g*(*c*) of composition where we omit the constant concentrations of all other components. In the following, we refer to function *g* as *concentration dependence*. Concentration buffering of the dilute phase concentration *c*_1_ occurs when *g* is insensitive to changes in *c*. To quantify concentration buffering, we define the *buffering strength* in the dilute phase

**Figure 1.**
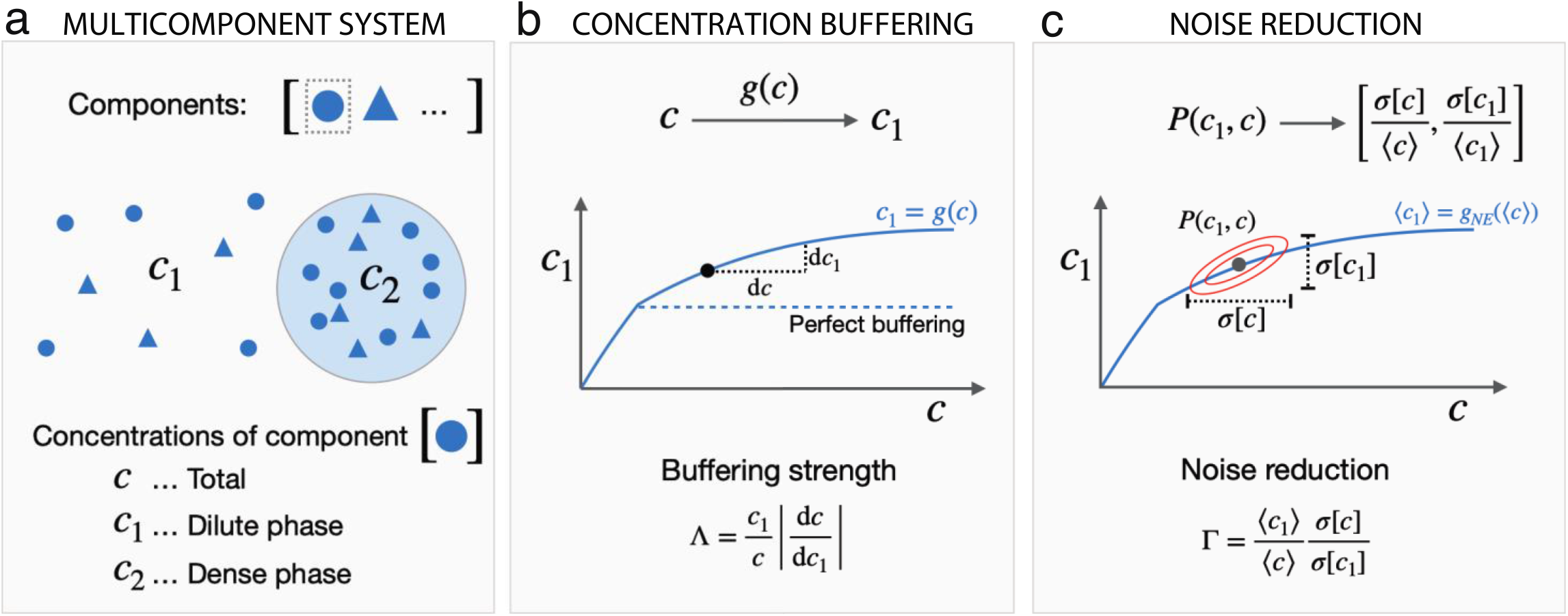
Concentration buffering and noise reduction in phase separating systems. **(a)** Scheme of a multicomponent system with two coexisting phases. Symbols *c*, c_1_ and *c*_2_ denote total-, dilute- and dense phase concentrations of the considered component *A*. **(b)** Concentration buffering in the dilute phase can be quantified by the buffering strength Λ, which we define as the inverse relative sensitivity of the dilute phase concentration *c*_1_ = *g*(*c*) with respect to changes in total concentration *c*, or equivalently, Λ = *d*Λ*og* c/*d*Λ*og* c_1_ **(c)** Noise reduction *Γ* can be defined as the ratio of the relative variabilities *σ*[. ]/⟨. ⟩ in total- and dilute phase concentration.

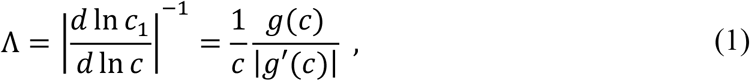

where *g′* denotes the derivative of *g* with respect to *c* (**Fig. 1b**). The buffering strength measures the logarithmic sensitivity of the dilute phase concentration *c*_1_ with respect to changes in total concentration *c*. We take the inverse to ensure that low sensitivities correspond to large buffering strengths. The logarithms imply that relative changes are considered. In a binary mixture at equilibrium with segregated macroscopic phases, *c*_1_ is constant when *c* increases, i.e., *g′* = 0 and the buffering strength Λ formally diverges. In this singular case, *c*_1_ is perfectly buffered as it is constrained to a fixed saturation concentration *c*_*sat*_ in the two-phase regime. In general, however, the buffering strength is a finite positive number, capturing how strongly *c*_1_ varies with *c*. As an example, a buffering strength of Λ=10 implies that when *c* changes by ∼10%, then *c*_1_ changes by only ∼1%. Even though the saturation concentration in this system is not fixed, the dilute phase concentration *c*_1_ is buffered effectively. The buffering strength can be generalized to non-equilibrium steady states where *g* becomes a non-equilibrium concentration dependence *g*_*NE*_. As we will discuss below, concentration buffering is often reduced in non-equilibrium conditions.

### Noise reduction

We now consider a phase separating system exhibiting concentration fluctuations, which we refer to as *noise*. The presence of noise renders both quantities, the total concentration *c* and the dilute phase concentration *c*_1_ time-dependent. Moreover, the relationship between *c* and *c*_1_ becomes stochastic as captured by the joint probability distribution *P*(*c*_1_, c). In the following, we consider the case where the statistics of the noise are stationary. In order to quantify noise reduction, we compare the probability distribution over total concentration *P*(*c*) = ∫ *P*(*c*_1_, c)*dc*_1_ to the probability distribution over dilute phase concentration *P*(*c*_1_) = ∫ *P*(*c*_1_, c)*dc* (**Fig. 1c**). A common dimensionless measure for the magnitude of noise is the coefficient of variation η[*x*] = *σ*[*x*]/⟨*x*⟩, where *σ*[*x*] and ⟨*x*⟩ are the standard deviation and mean of random variable *x*, respectively. The coefficient of variation measures the variability of *x* relative to its mean, which is useful when comparing quantities of different magnitude. Noise reduction in the dilute phase can then be defined as the ratio of coefficients of variation of total- and dilute phase concentration

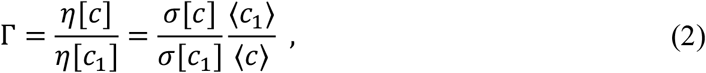

such that noise is reduced when Γ > 1 and η[*c*_1_] < η[*c*].

Molecular noise in cells is dynamic and spans a spectrum of timescales, ranging over orders of magnitude^14-16^. How fluctuations at different timescales are affected by a system depends on how the timescales of the noise compare to the intrinsic timescales of that system. As an example, a system may effectively suppress low-frequency fluctuations, but fail to suppress, or even amplify high-frequency fluctuations. Similar to the buffering strength,noise reduction is a quantitative property. In cells, a reduction of noise by a factor of as little as 1.5-2 can already be substantial and require high metabolic cost when realized through active biochemical feedback^17,18^.

### Concentration buffering and noise reduction are distinct concepts

Simple considerations suggest that concentration buffering and noise reduction may be closely related concepts^2,7,12,13^. This viewpoint originates from the idea that when *c*_1_ is insensitive to *c* (perfect buffering, Λ → ∞), it should buffer variations in *c* and leave *c*_1_ more or less unaffected even if *c* fluctuates. Comparing concentration buffering and noise reduction more closely, however, reveals that they are different and cannot be used synonymously. In general, concentration buffering cannot be used to predict noise reduction, nor can it provide bounds on it as we will show below.ΛTo better understand this, it is useful to first address the question under what special conditions concentration buffering and noise reduction do become similar. Consider an ensemble of equilibrium systems, each being at the thermodynamic limit with concentration relationship *g*(*c*). The systems only differ in the total concentrations *c* which are randomly distributed according to a probability distribution *P*(*c*). Physically, this corresponds to an equilibrium ensemble or to a quasi-static system, where the composition changes slowly compared to the relaxation to equilibrium. Since *c* varies, also *c*_1_ varies across the ensemble according to some probability distribution *P*(*c*_1_). If we consider small variations around the average total concentration ⟨*c*⟩, the standard deviation of *c*_1_ becomes *σ*[*c*_1_] ≃ *σ*[*c*] |*g′*(⟨*c*⟩) |. Using this expression in the definition of noise reduction (2)yields

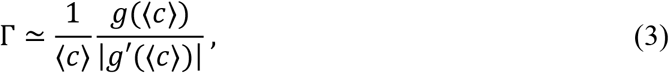

which is identical to the buffering strength (1) evaluated at the average total concentration ⟨*c*⟩. In other words, concentration buffering and noise reduction become similar for large equilibrium systems with slowly fluctuating composition. Note however, that this special case entails a strong simplification of the concept of noise reduction, essentially reducing it to the transformation of the random variable *c* by a static nonlinearity *g*.

In the cell biological context, molecular noise is discussed in the context of mesoscopic stochastic systems, whose degrees of freedom fluctuate dynamically and which are out of equilibrium. While noise reduction as defined in eq. (2) is well-defined for such systems, the buffering strength defined in eq. (1) needs to be reconsidered. This is because the total- and dilute phase concentrations are random variables, whose relationship cannot be expressed by a deterministic function. Alternatively, we can define the relationship between average concentrations ⟨*c*_1_⟩ = *g*_*NE*_(⟨*c*⟩) and use this function *g*_*NE*_ instead of *g* in the definition of the buffering strength in eq. (1). However, concentration buffering then becomes a property of average concentrations, while noise reduction is concerned with fluctuations around such averages. In general, these two quantities provide different information. In the following, we will use theory and numerical analyses to probe the relationship between concentration buffering and noise reduction in mesoscopic phase separating systems away from equilibrium.

### Stochastic dynamics of droplet systems in the presence of active synthesis and turnover of molecules

We consider a three-component mixture consisting of two solutes *A* and *B* and solvent *S*. While mixtures comprising more than three components could be handled analogously, a ternary mixture is sufficient for the following discussion. The copy numbers of the components are denoted by *a, b* and *s*, respectively. The solvent *S* is associated with a molecular volume *ν*, while the two solutes *A* and *B* are considered to have equal molecular volume *ν*_*A*_ = *ν*_*B*_ = *nν* for simplicity. We describe the composition of the mixture using volume fractions *ϕ* = *aν*_*A*_/*V* and ψ = *bν*_*B*_/*V*, which are particle concentrations multiplied by the respective molecular volumes *ν*_*A*_ and *ν*_*B*_. We denote the volume fractions of solute molecules in the dilute- and dense phase as *ϕ*_*α*_ = *ν*_*A*_*a*_*α*_/*V*_*α*_ and ψ_*α*_ = *ν*_*B*_*b*_*α*_/*V*_*α*_ for *α* ∈ 1, 2, respectively. Since volume fractions can be understood as rescaled concentrations, we will refer to them as concentrations in the following to simplify terminology. At equilibrium, this system can be described by a free energy

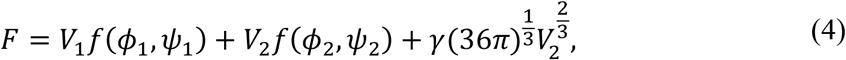

with *f* as a ternary Flory-Huggins free energy density, *γ* the surface tension, and *V*_1_ and *V*_2_ denoting the volumes of dilute and dense phase. The free energy density *f* is governed by three effective interaction parameters *X*_*AS*_, *X*_*BS*_ and *χ*_*AB*_ capturing effects of pair-wise interactions among *A*, B and *S*. Considering incompressibility (*V* = *V*_1_ + *V*_2_ = c*onst*. ) and conserved copy numbers *a* and *b*, the system can be described by three degrees of freedom, e.g., the dilute phase copy numbers *a*_1_, *b*_1_ and *s*_1_. The dependencies of dilute phase concentration on total concentration at equilibrium can be obtained by minimizing the free energy (4) with respect to *a*_1_, *b*_1_ and *s*_1_(**Supplementary text Section S.1.1**).

In the presence of molecular synthesis and degradation, the solute copy numbers *a* and *b* are no longer conserved. In this case, *a, b* and also *a*_1_, *b*_1_ and *s*_1_ are stochastic degrees of freedom. We describe copy number fluctuations of *a* and *b* using birth-and-death processes with fluctuating birth-rate^19^. In case of *a*, for instance, we consider a stochastic time-dependent birth-rate 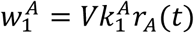 with rate constant 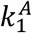 and a dimensionless stochastic process *r*_*A*_(*t*) . The process *r*_*A*_(*t*) can account for additional sources of randomness affecting the synthesis of components *A* such as fluctuations in the number of messenger RNA or extrinsic sources of variability^19,20^. We consider *r*_*A*_(*t*) to be at stationarity with mean ⟨*r*_*A*_⟩ and autocovariance function *κ*_*A*_(τ) = ⟨*r*_*A*_(*t*)*r*_*A*_(*t* + τ)⟩, where τ is the lag time. Degradation of molecules *A* takes place with rate 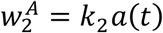, where 1/*K*_2_ is the average lifetime of a molecule. For simplicity, the same birth- and degradation rates are used in both phases. We can now calculate the mean and the variance of total concentration *ϕ* of solute *A*. The resulting stationary noise strength of total concentration *ϕ* reads

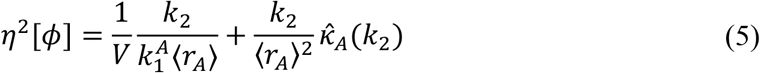

Where 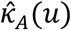 is the Laplace transform of the autocovariance function *κ*_*A*_(τ), which in eq. (5) is evaluated at the degradation rate *u* = *K*_2_ (**Supplementary text Section S.1.3**). Fluctuations of the second solute *B* are treated analogously.

To study how fluctuations in total concentration relate to fluctuations in the dilute (or dense) phase, we have to account for the kinetics of material exchange between phases. Specifically, we consider diffusion-limited exchange of solutes between dense- and dilute phase as well rapid relaxation of solvent to osmotic equilibrium. The kinetics of these events are governed by differences in chemical potentials, involving osmotic pressure and Laplace pressure (**Supplementary text Sections S.1.2 and S.1.4**). The statistical properties of this system can be described by a joint probability distribution *P*(*a*_1_, *b*_1_, s_1_, a, *b*, t), which can be described using a master equation formalism. Solving this equation, allows us to analyze both noise reduction and concentration buffering for components *A* and *B* in non-equilibrium conditions (**Supplementary text Sections S.1.4**).

### Non-equilibrium systems lack fixed saturation concentrations

We first consider concentration buffering and noise reduction in a simple binary mixture with only solute *A* and solvent *S* (no component *B*, ψ = 0) . Attractive interactions among solute molecules *A* or repulsive interactions between *A* and *S* are captured by a single interaction parameter *χ*_*AS*_ > 0. The total concentration *ϕ* of solutes *A* fluctuates due to synthesis and turnover as specified above. The stochastic process *r*_*A*_(*t*) is chosen to be a birth-and-death process with birth-rate λ_1_ and death-rate λ_2_*r*_*A*_(*t*) mimicking for example stochastic synthesis and degradation of messenger RNA. The stationary mean and autocovariance function of *r*_*A*_(*t*) are then given by 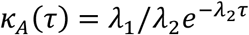.

To study concentration buffering in this system, we determine the dependence of the average dilute phase concentration on total concentration, ⟨*ϕ*_1_⟩ = *g*_*NE*_(⟨*ϕ*⟩) and compare it to the dilute phase concentration at equilibrium *ϕ*_1_ = *g*(*ϕ*). For a phase separated two-component system at equilibrium, we have *g*(*ϕ*) ≃ *ϕ*_*sat*_, where *ϕ*_*sat*_ is the equilibrium saturation concentration in the thermodynamic limit (**Fig. 2a**, black line). This is no longer the case in non-equilibrium conditions, where ⟨*ϕ*_1_⟩ = *g*_*NE*_(⟨*ϕ*⟩) increases with average total concentration ⟨*ϕ*⟩ (**Fig. 2a**, blue line, **Supplementary Fig. S1**). For the considered two-component system, approximate expressions can be derived if dilute phase concentrations *ϕ*_1_ and droplet volumes *V*_2_ are small (*dilute approximation*, see **Supplementary text Section S.1.5**). In the two-phase regime, *g*_*NE*_(⟨*ϕ*⟩) is given by

**Figure 2.**
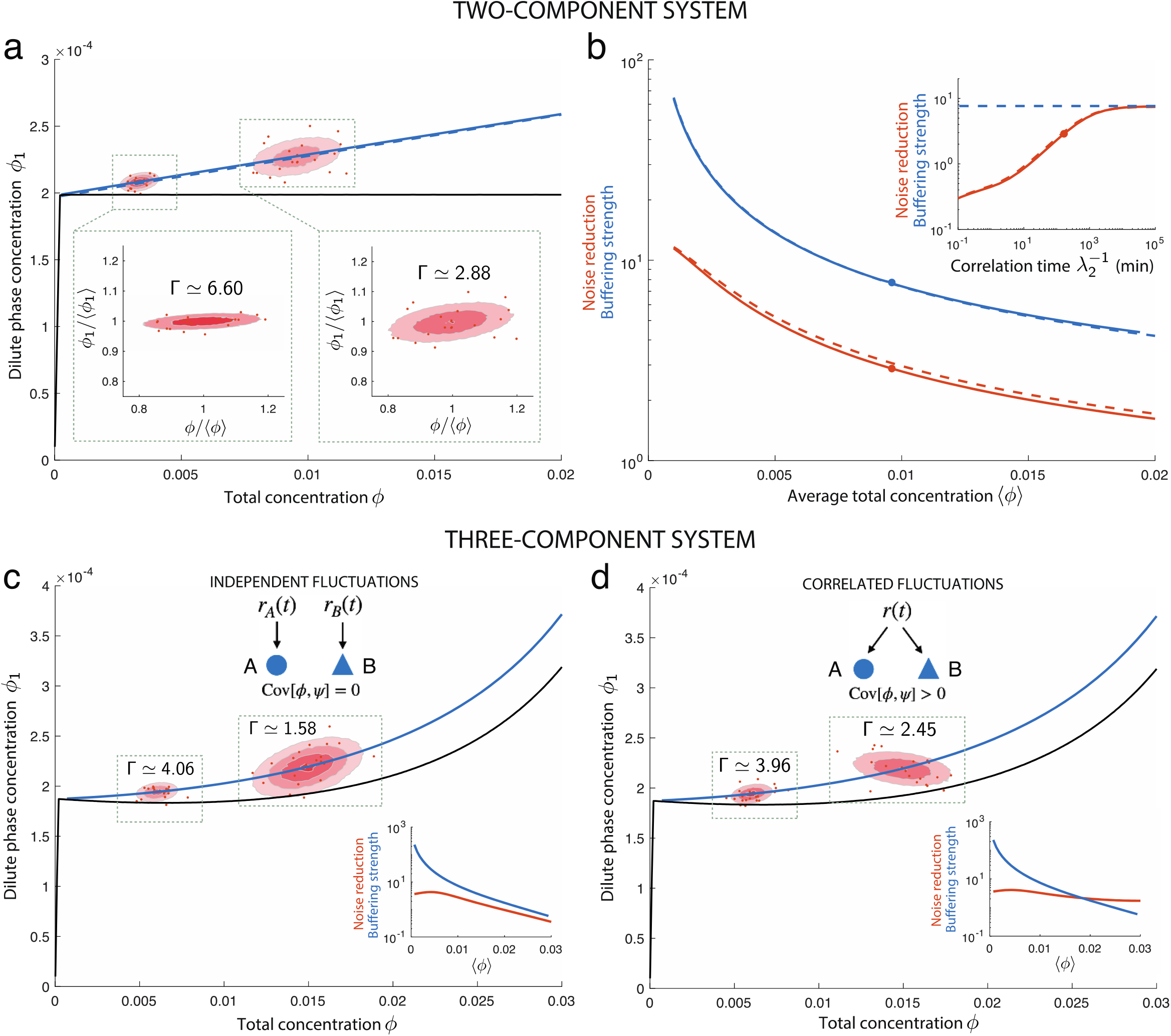
Concentration buffering and noise reduction out of equilibrium. **(a, b)** Two-component system consisting of a solute *A* and solvent *S*. The total concentration of *A* is described by *ϕ* = c*ν*_*A*_. **(a)** Dilute phase concentration *ϕ*_1_ as a function of total concentration *ϕ* of solute *A*. Joint distributions over total- and dilute phase concentrations are illustrated as ellipses for two different average total concentrations. Representative samples are shown as red dots. Insets show distributions over relative concentrations *ϕ*/⟨*ϕ*⟩ and *ϕ*_1_/⟨*ϕ*_1_⟩, illustrating relative variability of *ϕ* and *ϕ*_1_, respectively. **(b)** Comparison between buffering strength (blue) and noise reduction (red). The buffering strength and noise reduction under the dilute approximation are shown as dashed lines. Inset: buffering strength and noise reduction for varying correlation times 1/λ_2_ of the process *r*_*A*_(*t*). **(c, d)** Three-component system consisting of solutes *A* and *B* and solvent *S*. **(c)** Fluctuations in *A* and *B* are independent. **(d)** Fluctuations in *A* and *B* are correlated. Insets: buffering strength and noise reduction for varying ⟨*ϕ*⟩. Parameter values are provided in **Supplementary Tables S.1-S.3**.

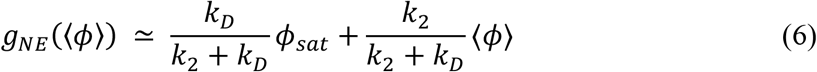

where *K*_*D*_ ∼ *D*/*V*^2/3^ is a kinetic coefficient that captures the diffusion-limited exchange of molecules between dilute- and dense phase with diffusion constant *D*. This demonstrates that the slope of *g*_*NE*_(⟨*ϕ*⟩) depends on the relative timescale between molecular turnover and partitioning, whereas perfect concentration buffering (⟨*ϕ*_1_⟩ → *ϕ*_*sat*_) is reached only when *K*_*D*_ is much larger than the degradation rate *K*_2_ and the phases reach equilibrium. At finite degradation rate, the two phases cannot equilibrate during the characteristic time of diffusion. Because the dilute phase concentration is not fixed, the apparent partition coefficient, which we define as ρ = ⟨*ϕ*_2_⟩/⟨*ϕ*_1_⟩ is also not constant but decreases with increasing ⟨*ϕ*⟩, i.e.,

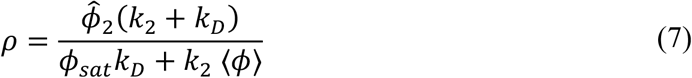

ΛWhere 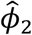 denotes the average dense phase concentration that is approximately constant (**Supplementary text Section S.1.5**). Thus, while a fixed saturation concentration and partition coefficient are hallmarks of binary systems at equilibrium^7,12,21^, these features are lost in non-equilibrium conditions where material is subject to production and turnover.

To compare these results to experiments, we used previously published data on the synthetic model protein 2NT-DDX4^YFP^, consisting of a fluorescent reporter fused to a disordered N-terminal domain^11,22^. The 2NT-DDX4^YFP^ construct was overexpressed in cells, which generated large variability in expression levels below and above the phase separation threshold. Condensates began to form at around 7 μ*M* but interestingly, average dilute phase concentrations did not remain constant but tended to increase with average total concentration (**Supplementary Fig. S2a**). Fitting equation (6) to the data, revealed a slope of *K*_2_/(*K*_2_ + *K*_*D*_) ≃ 0.04, corresponding to a case where protein partitioning is around 20 times faster than turnover (**Supplementary text Section S.2.1)**. We further display the partition coefficient for a smaller dataset from the same study^11^, where both dilute- and dense phase concentrations were measured, revealing that the partition coefficient decreases for increasing dilute phase concentration and hence, total concentration (**Supplementary Fig. S2a**, inset). The fit of the nonequilibrium binary model to the data show that the variable saturation concentration and partition coefficient are consistent even with a binary system with molecular synthesis and turnover. Therefore, from observed variable saturation concentrations or partition coefficients in a cell, one cannot infer that heterotypic interactions contribute to condensate formation, in contrast to previous suggestions^5,7,21^.

### A fixed saturation concentration is not required for effective concentration buffering and noise reduction

We next analyzed how the absence of a fixed saturation concentration affects concentration buffering and the reduction of noise. To this end, we determined the buffering strength and noise reduction of the non-equilibrium binary system for different average total concentrations ⟨*ϕ*⟩ (**Fig. 2a, b**). Both the buffering strength and noise reduction decrease with increasing average total concentration ⟨*ϕ*⟩, ranging from about 100-to 5-fold and 10-to 2-fold, respectively (**Fig. 2b**, blue and red lines). This demonstrates that both quantities depend strongly on the systems’ setpoint ⟨*ϕ*⟩ and that they can attain very large values even when the average dilute phase concentrations ⟨*ϕ*_1_⟩ varies strongly.

This analysis further reveals that the buffering strength and noise reduction can assume substantially different values for a given average total concentration ⟨*ϕ*⟩ (**Fig. 2b**). To better understand this difference, we make use of the dilute approximation (**Supplementary text Section S.1.5**), for which the buffering strength is

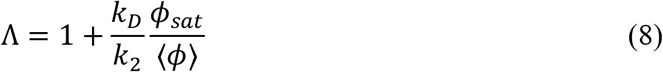

Noise reduction, by contrast takes a more complex form Γ = η[*ϕ*]/η[*ϕ*_1_] with 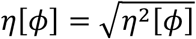 given by eq. (5) and

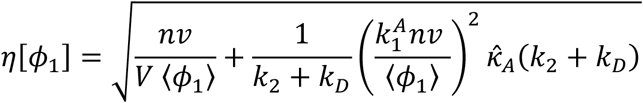

with ⟨*ϕ*_1_⟩ = *g*_*NE*_(⟨*ϕ*⟩) as defined in (6) (**Supplementary text Section S.1.5**). Comparing the buffering strength with noise reduction, we realize that the latter depends on the correlation function 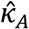 and the system size *V*, while the former does not. The size-dependence of noise reduction originates from intrinsic fluctuations of solute partitioning, synthesis and turnover. The dependence on 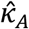 captures how temporal fluctuations of the birth-rate *r*_*A*_ propagate to *ϕ* and *ϕ*_1_, respectively. Notably, noise reduction becomes identical to the buffering strength in the limit of both large systems (*V* → ∞) and long correlation time of the birth-rate *r*_*A*_ (λ_2_ → 0) while keeping *K*_*D*_ fixed (**Supplementary text Section S.1.6**). However, cells do not typically operate close to these limits^14,20,23-25^. As an example, if we consider *r*_*A*_ to have a correlation time of 1/λ_2_ ≃ 3*h* (e.g., due to mRNA decay) together with a solute lifetime of 1/*K*_2_ ≃ 14*h* and a partitioning time of 1/*K*_*D*_ ≃ 2.6*min*, then the buffering strength differs from noise reduction by more than two-fold (**Fig. 2b**, inset, red dot). In summary, concentration buffering and noise reduction are distinct concepts, which become similar only in the extreme case of a very large and slowly varying system.

### Multicomponent systems can effectively buffer concentration and reduce noise

To generalize our analysis, we consider a three-component system, with both solutes *A* and *B* being present. For simplicity, we focus on the case of a purely heterotypic system, where *X*_*AS*_ = *X*_*BS*_ = 0 and *χ*_*AB*_ < 0. The stochastic processes *r*_*A*_ and *r*_*B*_ are considered to be independent but identical birth-and-death processes with mean ⟨*r*⟩ = λ_1_/λ_2_ and autocovariance function 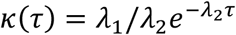as before. In the following, we focus on noise reduction and concentration buffering of solute *A*, but equivalent analyses apply to the second solute *B*. In line with previous observations^2,5,7^, this system displays a more complex concentration dependence, such that even at equilibrium, the average dilute phase concentration changes substantially with average total concentration (**Fig. 2c**, black line). The concentration dependence is further affected by the presence of synthesis and turnover of solutes *A* and *B* (**Fig. 2c**, blue line). As in the binary case, the buffering strength varies across different ⟨*ϕ*⟩ but in our example takes large values overall. Also, noise is reduced for a broad range of ⟨*ϕ*⟩, even though ⟨*ϕ*_1_⟩ varies substantially. These results are in line with experimental measurements of nucleolar component NPM1^7,11^, where dilute phase NPM1 concentration increased substantially upon overexpression^6^, while endogenous labelling of NPM1 revealed that noise reduction occurred at the physiological setpoint^11^ (**Supplementary Fig. S2b)**.

We further considered the case, where fluctuations in components *A* and *B* are positively correlated (**Fig. 2d**). This is inspired by previous theoretical work showing that coexisting concentrations of macroscopic systems are buffered when the total concentrations *ϕ* and ψ vary quasi-statically along tie lines^2^. Tie lines connect the coexisting compositions in a phase diagram. When variations of total concentrations occur along a tie line, the coexisting compositions at equilibrium do not change, while they do change if variations are not along a tie line^2,5^. Our analysis extends this concept to mesoscopic non-equilibrium systems where *ϕ* and ψ fluctuate on finite timescales. To account for correlations between *ϕ* and ψ, we consider the case where the same birth-process drives the synthesis of both components *A* and *B*, i.e., *r*_*A*_ = *r*_*B*_ = *r* with mean and autocovariance function as defined above. This leads to a positive correlation between *ϕ* and ψ which can be aligned with a tie line. While the predicted average concentrations ⟨*ϕ*⟩ and ⟨*ϕ*_1_⟩ are not affected by the presence of correlations between *ϕ* and ψ (compare blue lines in **Fig. 2c** and **d**), we find fluctuations of dilute phase concentration to depend in a notable manner on such correlations. In our example, fluctuations in total concentration *ϕ* and dilute phase concentration *ϕ*_1_ are positively correlated for small ⟨*ϕ*⟩, similarly to the case where *ϕ* and ψ are uncorrelated (**Fig. 2d**). For larger ⟨*ϕ*⟩, however, the correlations between *ϕ* and *ϕ*_1_ become negative, meaning that at the level of individual realizations (e.g., cells in a population), fluctuations leading to larger *ϕ* are associated with smaller *ϕ*_1_. At the same time, the average concentrations show the opposite behavior and ⟨*ϕ*_1_⟩ increases with increasing ⟨*ϕ*⟩ (**Fig. 2d**). This difference in behavior arises because in the case of average concentrations, buffering is characterized by the response of ⟨*ϕ*_1_⟩ to changes in ⟨*ϕ*⟩, while noise in *ϕ*_1_ is governed by correlated fluctuations of several variables. The exact statistics of *ϕ*_1_ depend on the interplay between the thermodynamic and nonequilibrium features of the system. This demonstrates that buffering of averages, and the behavior of fluctuations around averages can show very different behaviors. Moreover, while the buffering strength was consistently larger than noise reduction in our previous examples (**Fig. 2b** and **c**, inset), this is no longer true when fluctuations in *ϕ* and ψ are correlated. In this example, noise reduction can be smaller or larger than the buffering strength, depending on the setpoint ⟨*ϕ*⟩ (**Fig 2d**, inset). This analysis further illustrates that concentration buffering and noise reduction provide different insights into phase separating systems and that in general, one cannot be inferred from the other.

### A fixed saturation concentration implies effective concentration buffering but not noise reduction

So far, we have demonstrated that a fixed saturation concentration is not necessary for noise reduction and concentration buffering to occur. We next analyzed whether a fixed saturation concentration guarantees noise reduction and concentration buffering. To do so, we considered an example of a two-component system with fast phase separation dynamics exhibiting a weak dilute phase dependence *g*_*NE*_ on total concentration, resulting in a large buffering strength (**Supplementary Fig. S3a, b**, blue lines). Moreover, the parameters were chosen such that noise in total concentration was smaller than in the examples discussed above where noise reduction had been significant. In this case, even though partitioning is very fast on the timescale of solute turnover, noise reduction fails (**Supplementary Fig. S3a**, inset and **Supplementary Fig. S3b**, blue line). This is because noise reduction relies on partitioning noise to be weaker than the non-equilibrium fluctuations arising from protein production and degradation. As we have shown previously, partitioning noise therefore forms a lower bound on noise in the considered phase separating systems^11^. In summary, while a fixed saturation concentration ensures effective concentration buffering, it is not sufficient for noise reduction to take place.

## Conclusion

Using a series of examples, we have clarified the relationship between concentration buffering and noise reduction by phase separated compartments. We show that these two concepts are different and become similar only for large systems near thermodynamic equilibrium, i.e., if the composition varies slowly. In this special case, noise reduction can be related to concentration buffering governed by equilibrium phase diagrams^2,13^. As we have shown, however, these concepts can be substantially different when non-equilibrium fluctuations have a finite correlation time and when the finite system size is taken into account. In general, concentration buffering can neither predict noise reduction, nor can it provide bounds on it.

Inspired by previous discussions^7,12,13^, we have tested to what extent a fixed saturation concentration is important for noise reduction. We have shown that two-component and multicomponent systems with variable saturation concentrations can nevertheless reduce noise effectively. Conversely, we have shown that systems exhibiting a fixed saturation concentration may fail to suppress noise or may even increase it. Thus, a fixed saturation concentration is neither necessary nor sufficient for effective noise reduction to take place. If and to what extent a phase separated compartment reduces noise depends on the interplay between the thermodynamic and kinetic properties of that compartment.

Our results have broader implications for the analysis of intracellular condensates. As an example, it has been suggested that when the titration of a component leads to a change in dilute phase concentration, or the partition coefficient, this points towards heterotypic interactions governing the formation of these condensates^5,7,21^. This argument is based on equilibrium phase diagrams of multicomponent mixtures. However, our analysis shows that in the presence of non-equilibrium production and turnover, variable dilute phase concentrations and variable partition coefficients arise generically, independent of the type of the underlying interactions. Since in cells, material is subject to production and turnover, inference of interactions based on equilibrium phase diagrams is thus unreliable.

An important challenge in the condensate field is to understand how far traditional concepts from condensed matter physics can take us in studying intracellular compartments and where new concepts may be needed. Moving from simpler systems comprising few components to systems with larger compositional complexity is an important goal in this regard^2,7,21^. At the same time, we have to account for the dynamic complexity of cellular systems arising from stochastic, non-equilibrium processes^11,14,20,23^. Developing theoretical and experimental approaches to study noise and information processing in non-equilibrium multicomponent condensates will remain an important challenge in the future.

## Acknowledgements

We are grateful to David Zwicker, Adam Klosin, Akshaye Pal, Tyler Harmon, Alf Honigmann, Titus Franzmann and Anthony Hyman for critical discussion and feedback on this work. The authors thank the MPI-CBG, the MPI-PKS and the Max Planck Society for supporting this work. CZ acknowledges funding from the Deutsche Forschungsgemeinschaft (DFG, German Research Foundation) (ZE1216/1-1).

## Data availability

All data shown were published previously^7,11^. Source codes underlying this work are publicly available at https://www.github.com/zechnerlab/bufferingvsnoise.

## Supplementary Figures

**Figure S1.**
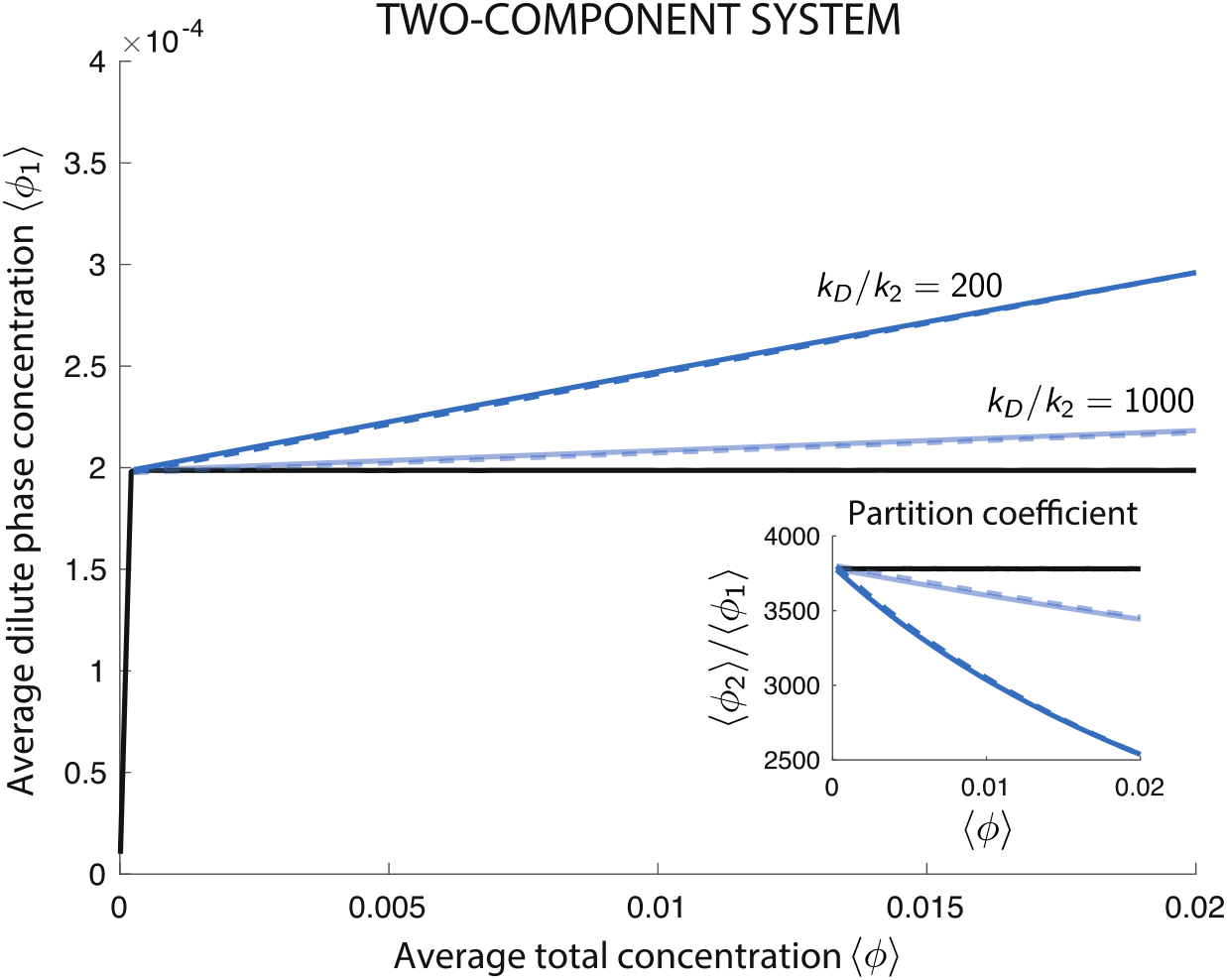
Concentration buffering in two-component systems with non-equilibrium synthesis and turnover. Concentration dependencies are shown for two different timescale ratios *K*_*D*_/*K*_2_ = 200 (dark blue) and *K*_*D*_/*K*_2_ = 1000 (light blue) and compared to equilibrium theory (black). Concentration dependencies obtained from the dilute approximation (**Supplementary text Section S.1.5**) are shown as dashed lines. Inset: corresponding apparent partition coefficients ρ = ⟨*ϕ*_2_⟩/⟨*ϕ*_1_⟩. Parameter values are provided in **Supplementary Table S.4**.

**Figure S2.**
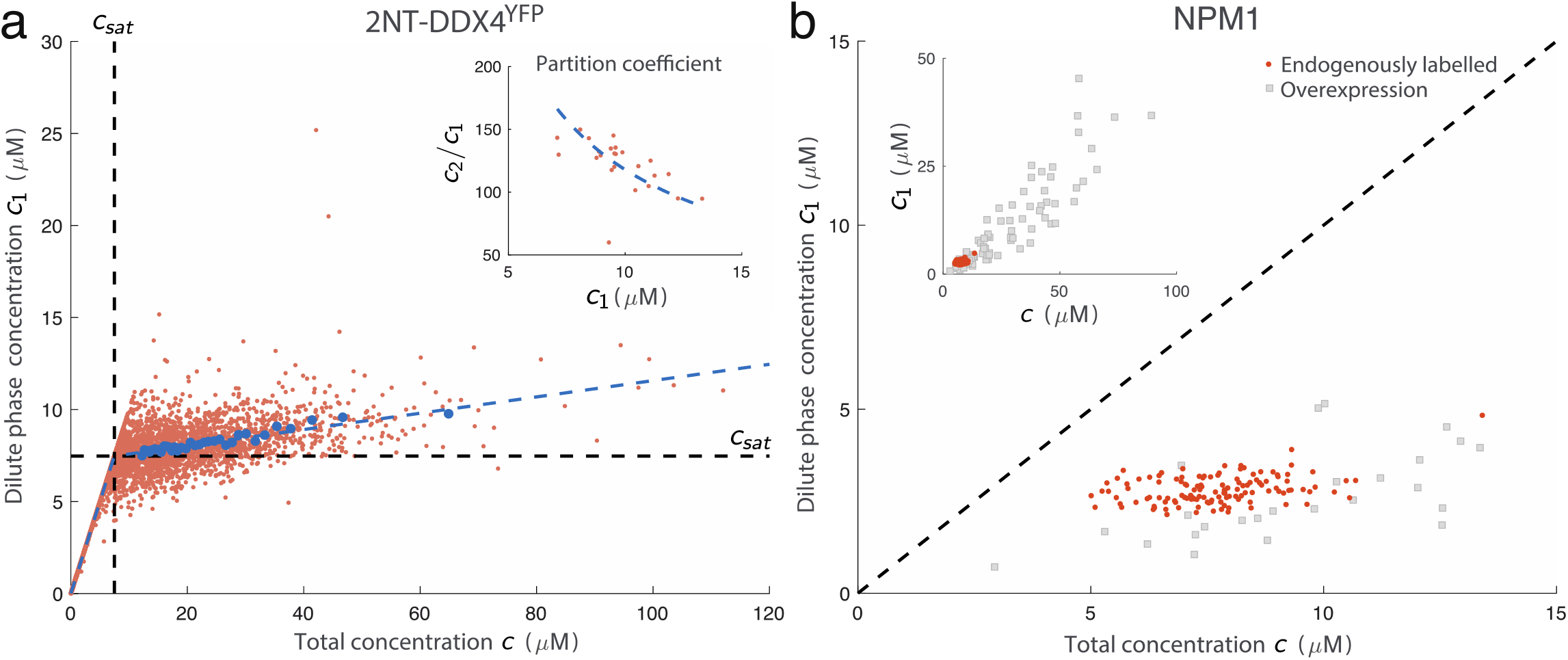
Experimental concentration measurements in synthetic and endogenous systems^7,11^. **(a)** Experimental measurements of 2NT-DDX4^YFP^ concentration reproduced from Klosin et al.^11^. Dilute phase-over total concentration of transiently expressed 2NT-DDX4^YFP^ (red dots). Average dilute- and total concentrations were estimated by binning cells into intervals of increasing total concentration and calculating averages in each subinterval (blue dots). Only cells with a total concentration larger than 12 μ*M* were used to select cells well within the two-phase regime. Eq. (6) for a two-component non-equilibrium system was fitted (blue dashed line) to the resulting concentration dependence, yielding estimates of the timescale ratio *K*_*D*_/*K*_2_ ≈ 21.569 ± 1.426 and the threshold concentration *c*_*sat*_ ≈ (7.471 ± 0.053) μ*M* (**Supplementary text Section S.2.1**). Inset: partition coefficient over dilute phase concentration in single cells (red dots). The dashed blue line shows the dilute approximation of the apparent partition coefficient ρ = ĉ_2_/⟨*c*_1_⟩ consistent with a two-component system with active synthesis and turnover (**Supplementary text Section S.1.5**). The average dense phase concentration ĉ_2_ was determined by averaging dense phase concentration over all cells. **(b)** Experimental measurements of NPM1 concentration reproduced from refs^7,11^. Endogenously labelled NPM1 intensity measurements from Klosin et al.^11^ (red dots) were rescaled to achieve a mean total concentration of 7.7 μ*M* as quantified previously^26^. Noise reduction was estimated from the data points as *Γ* ≃ 1.273 ± 0.103 using bootstrapping. The data from Riback et al.^7^ was plotted as published (grey squares). Inset: comparison between endogenous (red dots) and overexpressed (grey squares) NPM1 concentration.

**Figure S3.**
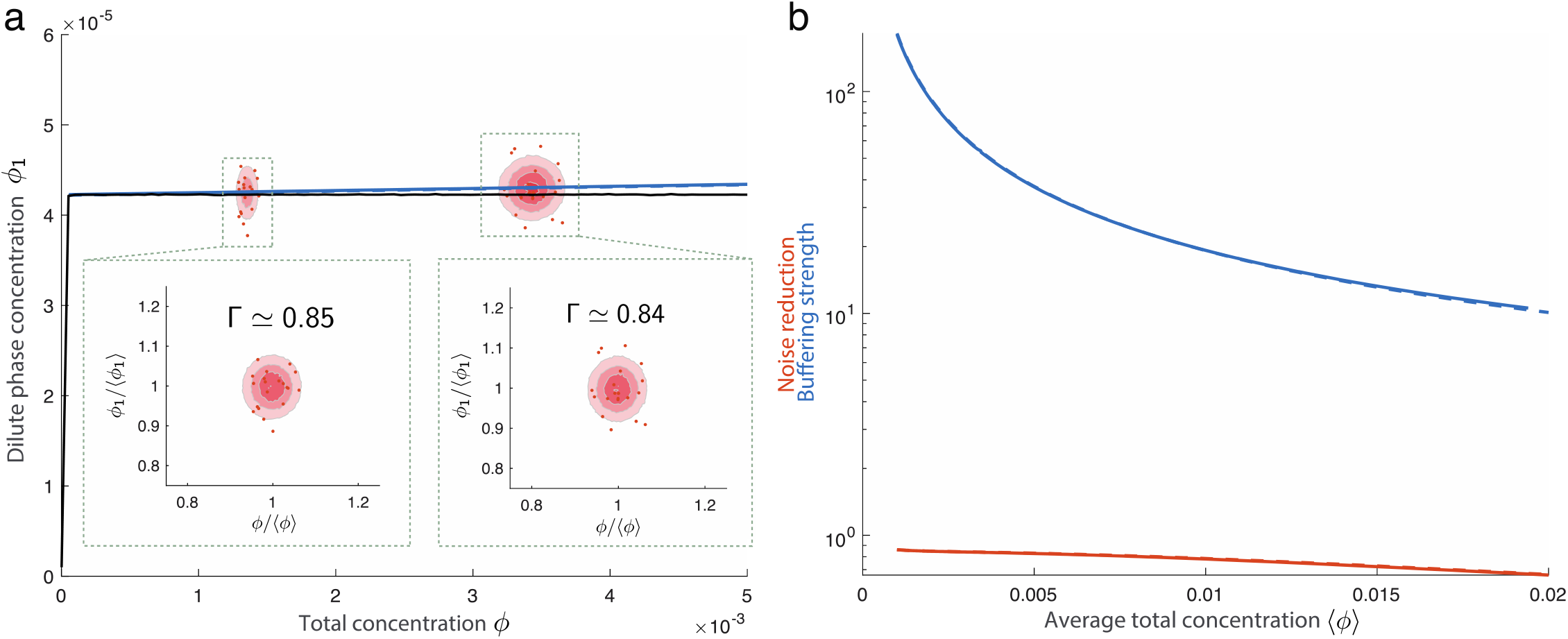
Concentration buffering and noise reduction in the presence of a nearly constant saturation concentration. **(a)** Scheme of a binary mixture where solute *A* is subject to active synthesis and turnover. **(b)** Average concentration and noise for varying average total concentration ⟨*ϕ*⟩ for solute *A*. Joint distributions over total- and dilute phase concentrations are illustrated as ellipses for two different average total concentrations. Representative samples are shown as red dots. Insets show distributions over relative concentrations *ϕ*/⟨*ϕ*⟩ and *ϕ*_1_/⟨*ϕ*_1_⟩, illustrating relative variability of *ϕ* and *ϕ*_1_, respectively. **(c)** Comparison between buffering strength (blue) and noise reduction (red). The buffering strength and noise reduction under the dilute approximation are shown as dashed lines. Parameter values are provided in **Supplementary Table S.5**.

## References

1 Banani, S. F., Lee, H. O., Hyman, A. A. & Rosen, M. K. Biomolecular condensates: organizers of cellular biochemistry. Nat Rev Mol Cell Bio 18, 285–298 (2017).

2 Deiyvri, D. & Safran, S. A. Physical theory of biological noise buffering by multicomponent phase separation. Proc Natl Acad Sci U S A 118, doi:10.1073/pnas.2100099118 (2021).

3 Jacobs, W. M. & Frenkel, D. Phase Transitions in Biological Systems with Many Components. Biophys J 112, 683–691, doi:10.1016/j.bpj.2016.10.043 (2017).

4 Jacobs, W. M. & Frenkel, D. Predicting phase behavior in multicomponent mixtures. J Chem Phys 139, 024108, doi:10.1063/1.4812461 (2013).

5 Choi, J. M., Dar, F. & Pappu, R. V. LASSI: A lattice model for simulating phase transitions of multivalent proteins. Plos Comput Biol 15, e1007028, doi:10.1371/journal.pcbi.1007028 (2019).

6 Mao, S., Kuldinow, D., Haataja, M. P. & Kosmrlj, A. Phase behavior and morphology of multicomponent liquid mixtures. Soft Matter 15, 1297–1311, doi:10.1039/c8sm02045k (2019).

7 Riback, J. A. et al. Composition-dependent thermodynamics of intracellular phase separation. Nature 581, 209–214, doi:10.1038/s41586-020-2256-2 (2020).

8 Graf, I. R. & Machta, B. B. Thermodynamic stability and critical points in multicomponent mixtures with structured interactions. Physical Review Research 4, 033144 (2022).

9 Holehouse, A. S. & Pappu, R. V. Functional Implications of Intracellular Phase Transitions. Biochemistry-Us 57, 2415–2423 (2018).

10 Stoeger, T., Battich, N. & Pelkmans, L. Passive Noise Filtering by Cellular Compartmentalization. Cell 164, 1151–1161 (2016).

11 Klosin, A. et al. Phase separation provides a mechanism to reduce noise in cells. Science 367, 464–468 (2020).

12 Chattaraj, A., Blinov, M. L. & Loew, L. M. The solubility product extends the buffering concept to heterotypic biomolecular condensates. Elife 10, doi:10.7554/eLife.67176 (2021).

13 Riback, J. A. & Brangwynne, C. P. Can phase separation buffer cellular noise? Science 367, 364–365, doi:10.1126/science.aba0446 (2020).

14 Rosenfeld, N., Young, J. W., Alon, U., Swain, P. S. & Elowitz, M. B. Gene regulation at the single-cell level. Science 307, 1962–1965, doi:10.1126/science.1106914 (2005).

15 Suter, D. M. et al. Mammalian genes are transcribed with widely different bursting kinetics. Science 332, 472–474, doi:10.1126/science.1198817 (2011).

16 Larson, D. R., Zenklusen, D., Wu, B., Chao, J. A. & Singer, R. H. Real-time observation of transcription initiation and elongation on an endogenous yeast gene. Science 332, 475–478, doi:10.1126/science.1202142 (2011).

17 Lestas, I., Vinnicombe, G. & Paulsson, J. Fundamental limits on the suppression of molecular fluctuations. Nature 467, 174–178 (2010).

18 Sartori, P. & Tu, Y. Free energy cost of reducing noise while maintaining a high sensitivity. Phys Rev Lett 115, 118102, doi:10.1103/PhysRevLett.115.118102 (2015).

19 Paulsson, J. Summing up the noise in gene networks. Nature 427, 415–418 (2004).

20 Elowitz, M. B., Levine, A. J., Siggia, E. D. & Swain, P. S. Stochastic gene expression in a single cell. Science 297, 1183–1186, doi:10.1126/science.1070919 (2002).

21 Mittag, T. & Pappu, R. V. A conceptual framework for understanding phase separation and addressing open questions and challenges. Mol Cell 82, 2201–2214, doi:10.1016/j.molcel.2022.05.018 (2022).

22 Nott, T. J. et al. Phase transition of a disordered nuage protein generates environmentally responsive membraneless organelles. Mol Cell 57, 936–947, doi:10.1016/j.molcel.2015.01.013 (2015).

23 Battle, C. et al. Broken detailed balance at mesoscopic scales in active biological systems. Science 352, 604–607, doi:10.1126/science.aac8167 (2016).

24 Yan, V. T., Narayanan, A., Wiegand, T., Julicher, F. & Grill, S. W. A condensate dynamic instability orchestrates actomyosin cortex activation. Nature 609, 597–604, doi:10.1038/s41586-022-05084-3 (2022).

25 Taniguchi, Y. et al. Quantifying E-coli Proteome and Transcriptome with Single-Molecule Sensitivity in Single Cells. Science 329, 533–538 (2010).

26 Freibaum, B. D., Messing, J., Yang, P., Kim, H. J. & Taylor, J. P. High-fidelity reconstitution of stress granules and nucleoli in mammalian cellular lysate. J Cell Biol 220, doi:10.1083/jcb.202009079 (2021).

