## Supplementary text for "On the relationship between concentration buffering and noise reduction in phase separating systems"

**S.3 Tables** 12

### S.1 Ternary mixture with non-equilibrium solute production and turnover

#### S.1.1 Equilibrium theory

We consider a system comprising two solutes $A$ and $B$ and solvent $S$, which phase separates into two phases of volumes $V_1$ and $V_2$, where $V_2$ is a droplet. The free energy of this system is written as

$$F = V_1 f(\phi_1, \psi_1) + V_2 f(\phi_2, \psi_2) + \gamma \mathcal{A}_2, \tag{1}$$

with $\mathcal{A}_2 = (36\pi)^{1/3} V_2^{2/3}$ as the surface area of the droplet and $\gamma$ denoting surface tension. For the free energy density $f$ of a homogenous phase, we consider a ternary Flory-Huggins model

$$\begin{aligned} f(\phi, \psi) = \frac{k_B T}{v} \Big[ &\chi_{AS} \phi (1 - \phi - \psi) + \chi_{AB} \phi \psi + \chi_{BS} \psi (1 - \phi - \psi) \\ &+ \frac{\phi}{n} \log \phi + \frac{\psi}{n} \log \psi + (1 - \phi - \psi) \log (1 - \phi - \psi) \Big], \end{aligned} \tag{2}$$

with effective interaction parameters $\chi_{AS}$, $\chi_{AB}$ and $\chi_{BS}$. For simplicity, we consider the case where the solutes $A$ and $B$ have identical molecular volume $nv$ with $v$ denoting the molecular volume of solvent $S$ and $n$ as a positive number. We chose to describe the system using the degrees of freedom $a_1$, $b_1$ and $s_1$, corresponding to the copy numbers of $A$, $B$ and $S$ molecules in the dilute phase of volume $V_1$, respectively. We then have $\phi_1 = a_1 nv/V1$, $\psi_1 = b_1 nv/V1$, $V_1 = v(a_1 n + b_1 n + s_1)$ and we use $a_1 = a - a_1$, $b_2 = b - b_1$ and $V_2 = V - V_1$ due to particle number conservation ($a = a_1 + a_2$ and $b = b_1 + b_2$) and incompressibility ($V = V_1 + V_2 = const.$). In terms of the degrees of freedom $a_1$, $b_1$ and $s_1$, the free energy reads

$$
\begin{aligned}
F(a_1, b_1, s_1; a, b) = v(a_1 n + b_1 n + s_1) f\left(\frac{a_1 n}{a_1 n + b_1 n + s_1}, \frac{b_1 n}{a_1 n + b_1 n + s_1}\right) \\
+ \left[V - v(a_1 n + b_1 n + s_1)\right] f\left(\frac{(a - a_1) n v}{V - v(a_1 n + b_1 n + s_1)}, \frac{(b - b_1) n v}{V - v(a_1 n + b_1 n + s_1)}\right) \\
+ \gamma (36\pi)^{1/3} \left[V - v(a_1 n + b_1 n + s_1)\right]^{2/3}.
\end{aligned} \tag{3}
$$

At equilibrium, $a_1$, $b_1$ and $s_1$ exhibit Boltzmann statistics

$$
P_{EQ}(a_1, b_1, s_1 \mid a, b) = \frac{1}{Z} e^{-\frac{F(a_1, b_1, s_1; a, b)}{k_B T}}, \tag{4}
$$

where

$$
Z = \sum_{(a, b_1, s_1) \in \mathcal{S}} e^{-\frac{F(a_1, b_1, s_1)}{k_B T}} \tag{5}
$$

is the partition function and $\mathcal{S} = [0, a] \times [0, b] \times [0, V/v - n(a + b)]$. Equilibrium quantities can be determined by calculating moments of the Boltzmann distribution as a function of $a$ or $b$. In case of solute $A$, for instance, we define

$$
\langle \phi_1 \mid \phi, \psi \rangle = \langle \phi_1 \mid a, b \rangle = \sum_{(a, b_1, s_1) \in \mathcal{S}} \frac{a_1 n}{a_1 n + b_1 n + s_1} P_{EQ}(a_1, b_1, s_1 \mid a, b). \tag{6}
$$

In eq. (6), the symbol $\langle \phi_1 \mid \phi, \psi \rangle$ denotes the conditional expectation of $\phi_1$ given a particular $\phi$ and $\psi$. As we are interested in how the dilute phase concentration of one component (e.g., $\phi_1$) changes with total concentration of the same component (e.g., $\phi$) we will drop the dependency on total concentration of the other component (e.g., $\psi$) in our notation. In case of component $A$, for instance, we define

$$
g(\phi) = \langle \phi_1 \mid \phi, \psi \rangle \tag{7}
$$

as $\psi$ is fixed. For sufficiently abundant systems with approximately Gaussian fluctuations, the above expectation may be approximated by the minimum of the free energy, or equivalently, the mode of the underlying Boltzmann distribution and we will used this approximation when calculating equilibrium concentration dependencies. Numerically, the minimum of the free energy was determined using the *fmincon* function of MATLAB (version 9.10.0.1602886 (R2021a), The MathWorks Inc., Natick, Massachusetts), using the interior-point algorithm under default settings(`https://github.com/zechnerlab/bufferingvsnoise`).

#### S.1.2 Kinetics of solute and solvent partitioning

To study concentration fluctuations in non-equilibrium conditions (see Section S.1.3), we require a kinetic description of the considered phase separating system. To this end, we consider three pairs of molecular exchange events

$$
(a_1, b_1, s_1) \underset{w_A^+}{\overset{w_A^-}{\rightleftharpoons}} (a_1 - 1, b_1, s_1 + n) \tag{8}
$$

$$
(a_1, b_1, s_1) \underset{w_B^+}{\overset{w_B^-}{\rightleftharpoons}} (a_1, b_1 - 1, s_1 + n) \tag{9}
$$

$$
(a_1, b_1, s_1) \underset{w_S^+}{\overset{w_S^-}{\rightleftharpoons}} (a_1, b_1, s_1 - 1). \tag{10}
$$

The first two rows correspond to exchange of solutes $A$ and $B$ in and out of the dense phase, while the third row captures volume fluctuations of the phases due to solvent exchange. These events are driven by generalized thermodynamic forces. Thermodynamics requires the rates to obey the following conditions

$$\log \frac{w_A^-(a_1, b_1, s_1)}{w_A^+(a_1 - 1, b_1, s_1 + n)} = -\frac{1}{k_B T}\left[F(a_1 - 1, b_1, s_1 + n) - F(a_1, b_1, s_1)\right] \tag{11}$$

$$\log \frac{w_B^-(a_1, b_1, s_1)}{w_B^+(a_1, b_1 - 1, s_1 + n)} = -\frac{1}{k_B T}\left[F(a_1, b_1 - 1, s_1 + n) - F(a_1, b_1, s_1)\right] \tag{12}$$

$$\log \frac{w_S^-(a_1, b_1, s_1)}{w_S^+(a_1, b_1, s_1 - 1)} = -\frac{1}{k_B T}\left[F(a_1, b_1, s_1 - 1) - F(a_1, b_1, s_1)\right], \tag{13}$$

where $w_A^{+/-}$, $w_B^{+/-}$ and $w_S^{+/-}$ are event probabilities per unit time. Considering copy numbers to be sufficiently large, the free energy differences in (11-13) can be approximated as directional derivatives. More precisely, we approximate $F(a_1 + \nu_a, b_1 + \nu_b, s_1 + \nu_s) - F(a_1, b_1, s_1) \simeq (\nu_a \ \nu_b \ \nu_s)\nabla F(a_1, b_1, s_1)$, with $\nabla F(a_1, b_1, s_1) = (\partial_{a_1} \ \partial_{b_1} \ \partial_{s_1})^T F(a_1, b_1, s_1)$ as the gradient of the free energy. This leads to

$$\log \frac{w_A^-(a_1, b_1, s_1)}{w_A^+(a_1 - 1, b_1, s_1 + n)} = -\begin{pmatrix}-1 & 0 & n\end{pmatrix}\nabla F(a_1, b_1, s_1) = -\frac{nv}{k_B T}(\mu_2^A - \mu_1^A) \tag{14}$$

$$\log \frac{w_B^-(a_1, b_1, s_1)}{w_B^+(a_1, b_1 - 1, s_1 + n)} = -\begin{pmatrix}0 & -1 & n\end{pmatrix}\nabla F(a_1, b_1, s_1) = -\frac{nv}{k_B T}(\mu_2^B - \mu_1^B) \tag{15}$$

$$\log \frac{w_S^-(a_1, b_1, s_1)}{w_S^+(a_1, b_1, s_1 - 1)} = -\begin{pmatrix}0 & 0 & -1\end{pmatrix}\nabla F(a_1, b_1, s_1) = -\frac{v}{k_B T}\left[\Pi_2 - \Pi_1\right] - \frac{\gamma}{k_B T}\partial_{s_1}\mathcal{A}_2, \tag{16}$$

where $\mu_\alpha^A = \partial_{\phi_\alpha} f(\phi_\alpha, \psi_\alpha)$ and $\mu_\alpha^B = \partial_{\psi_\alpha} f(\phi_\alpha, \psi_\alpha)$ are the exchange chemical potential of component $A$ and $B$ in phase $\alpha \in \{1, 2\}$, $\Pi_2 - \Pi_1$ is the osmotic pressure difference with $\Pi_\alpha = f(\phi_\alpha, \psi_\alpha) - \phi\mu_\alpha^A - \psi\mu_\alpha^B$ and $\gamma\partial_{s1}\mathcal{A}_2/v$ is the Laplace pressure.

We choose the partitioning of solutes into the dense phase to be diffusion-limited with $w_A^+(a_1, b_1, s_1) = 6D/V_1^{2/3}a_1$ and $w_B^+(a_1, b_1, s_1) = 6D/V_1^{2/3}b_1$, where we consider the two solutes $A$ and $B$ to have the same diffusion constant $D$ for simplicity. The corresponding reverse rates $w_A^-$ and $w_B^-$ are then chosen to satisfy (14) and (15). Solvent exchange (10) is expected to be fast on the timescale of solute exchange, effectively keeping the system close to osmotic equilibrium. We therefore consider the limit where the kinetic coefficients associated with $w_S^+$ and $w_S^-$ are fast in comparison to solute exchange, effectively eliminating one kinetic mode of this system. Practically, this was achieved by choosing $w_S^-(a_1, b_1, s_1) = k_S$, determining the reverse rate $w_s^+$ to satisfy (14) and setting the kinetic coefficient $k_S$ to large values (see Tables S.1-S.5).

#### S.1.3 Accounting for non-equilibrium production and turnover of solutes

In the presence of noise, the total concentrations $\phi$ and $\psi$ are no longer conserved. To describe solute fluctuations, we use stochastic birth-and-death processes with fluctuating birth-rates. Despite their simplicity, these processes are remarkably consistent with concentration fluctuations in cells [1, 2, 3, 4]. We describe synthesis and turnover of a component $X \in \{A, B\}$ as

$$(x, s) \underset{w_{B,2}^d}{\overset{w_{B,2}^b}{\rightleftharpoons}} (x + 1, s - n), \tag{17}$$

where whenever a molecule $x$ is produced or degraded, $n$ solvent molecules are removed or added to the system, leaving the total volume conserved. We further set $w_X^b = Vk_1^X r_X(t)$ where $k_1$ is a birth rate per volume and $w_X^d = k_2 x(t)$, where for simplicity we consider the case where both solutes $A$ and $B$ are degraded with the same rate constant $k_2$. The stochastic processes $r_X(t)$ capture fluctuations in the synthesis rate, for instance due to transcriptional noise or extrinsic variability. We consider $r_X(t)$ to be a stationary stochastic process with mean $\langle r\rangle$ and autocovariance function $\kappa_X(\tau) = \langle r_X(t)r_X(t+\tau)\rangle - \langle r\rangle^2$ for $X \in \{A, B\}$. In the presence of phase coexistence, both synthesis and turnover events can take place in both phases $i = \{1, 2\}$ with rates $w^b_{X,i} = V_i k_1^X r_X(t)$ and $w^d_{X,i} = k_2 x_i(t)$, leaving the statistics of total concentration $\phi$ and $\psi$ unchanged. This choice is useful to compare fluctuations in the presence and absence of phase coexistence. Note that in order to avoid negative solvent copy numbers, we require $w^b_{X,i} = 0$ if $s_i < n$. However, since we consider regimes where solvent molecules are much more abundant than solute molecules, we neglect this constraint in our calculations. In the following, we will denote by $\mathcal{BD}(w^b_X, w^d_X)$ a birth-and-death process $X$ with birth-rate $w^b_X$ and death-rate $w^d_X$.

To derive the statistics of total solute concentration, we consider the birth-and-death process $\mathcal{BD}(Vk_1 r(t), k_2 x(t))$. For a particular realization of the process $r(t)$, the probability of having $x$ molecules at time $t$ can be described by a master equation

$$\begin{aligned}\frac{\mathrm{d}}{\mathrm{d}t}P(x,t) &= Vk_1 r(t)P(x-1,t) + k_2(x+1)P(x+1,t) \\ &\quad - \left[Vk_1 r(t) + k_2 x\right]P(x,t),\end{aligned} \tag{18}$$

with $P(x,t) := P(x(t) \mid r_0^t)$ as the copy number distribution of molecules $X$ at time $t$ given a realization $r_0^t$ of the process $r(t)$. The conditional mean and variance of $x$ are given by

$$\langle x(t) \mid r_0^t\rangle = \mathrm{Var}[x(t) \mid r_0^t] = Vk_1 \int_0^t e^{-k_2(t-\tau)} r(\tau)\mathrm{d}\tau \tag{19}$$

where we have chosen $\langle x(0) \mid r_0\rangle = \mathrm{Var}[x(0) \mid r_0] = 0$ for convenience (the initial conditions are irrelevant since we are interested in the long-term behavior). The stationary mean of $x(t)$ is readily obtained by taking the expectation of (19) and letting $t \to \infty$, which yields

$$\langle x\rangle = \frac{Vk_1\langle r\rangle}{k_2}. \tag{20}$$

For the variance of $x(t)$, we make use of the law of total variance, which states that

$$\mathrm{Var}[x(t)] = \langle \mathrm{Var}[x(t) \mid r_0^t]\rangle + \mathrm{Var}[\langle x(t) \mid r_0^t\rangle]. \tag{21}$$

The first term on the rhs of (21) is the same as the stationary mean, i.e., $\langle \mathrm{Var}[x(t) \mid r_0^t]\rangle \xrightarrow{t\to\infty} k_1 V\langle r\rangle/k_2$. The second term on the rhs of (21) is given by

$$\begin{aligned}\mathrm{Var}[\langle x(t) \mid r_0^t\rangle] &= \mathrm{Var}\left[Vk_1 \int_0^t e^{-k_2(t-\tau)} r(\tau)\mathrm{d}\tau\right] \\ &= k_1^2 V^2 e^{-2k_2 t}\mathrm{Var}\left[\int_0^t e^{k_2\tau} r(\tau)\mathrm{d}\tau\right].\end{aligned} \tag{22}$$

The variance of the integral can be rewritten as

$$\begin{aligned}\mathrm{Var}\left[\int_0^t e^{k_2\tau} r(\tau)\mathrm{d}\tau\right] &= \left\langle \left(\int_0^t e^{k_2\tau} r(\tau)\mathrm{d}\tau\right)^2 \right\rangle - \left(\int_0^t e^{k_2\tau}\langle r(\tau)\rangle\mathrm{d}\tau\right)^2 \\ &= \int_0^t\int_0^t e^{k_2(\tau+\tau')}\langle r(\tau)r(\tau')\rangle\mathrm{d}\tau\mathrm{d}\tau' - \int_0^t\int_0^t e^{k_2(\tau+\tau')}\langle r(\tau)\rangle\langle r(\tau')\rangle\mathrm{d}\tau\mathrm{d}\tau' \\ &= \int_0^t\int_0^t e^{k_2(\tau+\tau')}\kappa(\tau'-\tau)\mathrm{d}\tau\mathrm{d}\tau'.\end{aligned} \tag{23}$$

Using the fact that the autcovariance is time-symmetric, we can simplify this expression to

$$\begin{aligned}&\int_0^t\int_0^t e^{k_2(\tau+\tau')}\kappa(\tau'-\tau)\mathrm{d}\tau\mathrm{d}\tau' \\ &\quad = \int_0^t\int_{\tau'}^t e^{k_2(\tau+\tau')}\kappa(\tau'-\tau)\mathrm{d}\tau\mathrm{d}\tau' + \int_0^t\int_0^{\tau'} e^{k_2(\tau+\tau')}\kappa(\tau'-\tau)\mathrm{d}\tau\mathrm{d}\tau' \\ &\quad = \int_0^t\int_0^{\tau} e^{k_2(\tau+\tau')}\kappa(\tau-\tau')\mathrm{d}\tau'\mathrm{d}\tau + \int_0^t\int_0^{\tau'} e^{k_2(\tau+\tau')}\kappa(\tau'-\tau)\mathrm{d}\tau\mathrm{d}\tau' \\ &\quad = 2\int_0^t\int_0^{\tau'} e^{k_2(\tau+\tau')}\kappa(\tau'-\tau)\mathrm{d}\tau\mathrm{d}\tau'.\end{aligned} \tag{24}$$

The variance of the conditional mean is thus given by

$$\mathrm{Var}[\langle x(t) \mid r_0^t\rangle] = 2k_1^2 V^2 e^{-2k_2 t} \int_0^t \int_0^{\tau'} e^{k_2(\tau+\tau')} \kappa(\tau' - \tau) \mathrm{d}\tau \mathrm{d}\tau'. \tag{25}$$

We next calculate the Laplace transform of this equation, which will allow us to determine a general expression for $\mathrm{Var}[\langle x(t) \mid r_0^t\rangle]$ in the limit of $t \to \infty$. To this end, we first rewrite eq. (26) as

$$\mathrm{Var}[\langle x(t) \mid r_0^t\rangle] = 2k_1^2 V^2 e^{-2k_2 t} \int_0^t e^{k_2\tau'} \int_0^{\tau'} e^{k_2\tau} \kappa(\tau' - \tau) \mathrm{d}\tau \mathrm{d}\tau', \tag{26}$$

such that the inner integral now has the form of a convolution integral. To calculate the Laplace transform of eq. (26), we make use of the properties [5]

$$\mathcal{L}\left\{\int_0^t g(s) f(t-s) \mathrm{d}s\right\} = \hat{g}(u)\hat{f}(u) \tag{27}$$

$$\mathcal{L}\left\{e^{\beta t} g(t)\right\} = \hat{g}(u-\beta) \tag{28}$$

$$\mathcal{L}\left\{\int_0^t g(s) \mathrm{d}s\right\} = \frac{1}{u}\hat{g}(u), \tag{29}$$

where $\hat{g}(u) = \int_0^\infty g(t)e^{-ut}\mathrm{d}t$ and $\hat{f}(u) = \int_0^\infty f(t)e^{-ut}\mathrm{d}t$ are the one-sided Laplace transforms of functions $g$ and $f$, respectively and $\beta$ is a constant. Using these properties, we obtain

$$\mathcal{L}\left\{\mathrm{Var}[\langle x(t) \mid r_0^t\rangle]\right\} = \frac{2k_1^2 V^2}{u(u+2k_2)} \hat{\kappa}(u+k_2), \tag{30}$$

with $\hat{\kappa}(u) = \int_0^\infty \kappa(t)e^{-ut}\mathrm{d}t$ as the Laplace transform of the autocovariance function $\kappa$. Further using the property $\lim_{t\to\infty} g(t) = \lim_{u\to 0} u\hat{g}(u)$ [5], we obtain for the long-term variance of the conditional mean (assuming that it exists)

$$\lim_{t\to\infty} \mathrm{Var}[\langle x(t) \mid r_0^t\rangle] = \lim_{u\to 0} \frac{2k_1^2 V^2}{u+2k_2} \kappa(u+k_2) = V^2 \frac{k_1^2}{k_2} \hat{\kappa}(k_2). \tag{31}$$

In total, we obtain for the stationary variance of $x(t)$

$$\mathrm{Var}[x] = V\frac{k_1}{k_2}\langle r\rangle + V^2 \frac{k_1^2}{k_2}\hat{\kappa}(k_2). \tag{32}$$

The relative noise defined as the squared coefficient of variation $\eta^2[x] = \mathrm{Var}[x]/\langle x\rangle^2$ is given by

$$\eta^2[x] = \frac{1}{V}\frac{k_2}{k_1\langle r\rangle} + \frac{k_2}{\langle r\rangle^2}\hat{\kappa}(k_2). \tag{33}$$

The first term on the right-hand side of (33) can be identified as $1/\langle x\rangle$ and thus exhibits Poissonian scaling. The second term quantifies the amount of noise that propagates from $r(t)$ to $x$, which depends on the timescales of $r(t)$ through the autocovariance function $\kappa(\tau)$.

Note that the noise strength $\eta^2[x]$ is invariant upon rescaling of $x$ such that (33) applies also to the concentration (and volume fraction) of $X$. The noise strength of total concentrations $\phi$ and $\psi$ are thus given by

$$\eta^2[\phi] = \frac{1}{V}\frac{k_2}{k_1^A\langle r_A\rangle} + \frac{k_2}{\langle r_A\rangle^2}\hat{\kappa}_A(k_2) \tag{34}$$

$$\text{and}\quad \eta^2[\psi] = \frac{1}{V}\frac{k_2}{k_1^B\langle r_B\rangle} + \frac{k_2}{\langle r_B\rangle^2}\hat{\kappa}_B(k_2). \tag{35}$$

#### S.1.4 Droplet kinetics in the presence of non-equilibrium production and turnover of solutes

To study how non-equilibrium fluctuations in total concentration are affected by phase coexistence, we consider a system that accounts for solute and solvent partitioning as well as synthesis and degradation of molecules in each phase. The state of this system is characterized by seven degrees of freedom $(a_1(t), b_1(t), s_1(t), a(t), b(t), r_A(t), r_B(t))$, which evolve according to the events

$$(a_1, b_1, s_1, a, b) \underset{w_A^+}{\overset{w_A^-}{\rightleftharpoons}} (a_1 - 1, b_1, s_1 + n, a, b) \tag{36}$$

$$(a_1, b_1, s_1, a, b) \underset{w_B^+}{\overset{w_B^-}{\rightleftharpoons}} (a_1, b_1 - 1, s_1 + n, a, b) \tag{37}$$

$$(a_1, b_1, s_1, a, b) \underset{w_S^+}{\overset{w_S^-}{\rightleftharpoons}} (a_1, b_1, s_1 - 1, a, b) \tag{38}$$

$$(a_1, b_1, s_1, a, b) \underset{w_{A,1}^d}{\overset{w_{A,1}^b}{\rightleftharpoons}} (a_1 + 1, b_1, s_1 - n, a + 1, b) \tag{39}$$

$$(a_1, b_1, s_1, a, b) \underset{w_{A,2}^d}{\overset{w_{A,2}^b}{\rightleftharpoons}} (a_1, b_1, s_1, a + 1, b) \tag{40}$$

$$(a_1, b_1, s_1, a, b) \underset{w_{B,1}^d}{\overset{w_{B,1}^b}{\rightleftharpoons}} (a_1, b_1 + 1, s_1 - n, a + 1, b) \tag{41}$$

$$(a_1, b_1, s_1, a, b) \underset{w_{B,2}^d}{\overset{w_{B,2}^b}{\rightleftharpoons}} (a_1, b_1, s_1, a, b + 1), \tag{42}$$

with rates as defined above. The birth-processes $r_A(t)$ and $r_B(t)$ have so far not been specified beyond their first- and second-order statistics. This system can be described by a probability distribution $P(a_1, b_1, s_1, a, b, r_A, r_B, t)$, which satisfies a master equation. To study fluctuations in total- and dilute phase volume fractions we calculated the first- and second-order moments of the degrees of freedom. Since the system is nonlinear, however, we cannot calculate the moments exactly. Similarly to our previous work [6], we address this problem by employing the linear noise approximation (LNA) [7], which yields first- and second order moments consistent with a Gaussian approximation. The LNA is valid for large systems consisting of sufficiently many particles. To apply the LNA to the considered system, we describe both $r_A(t)$ and $r_B(t)$ as birth-and-death processes $\mathcal{BD}(\lambda_1, \lambda_2 r_X(t))$ for $X \in \{A, B\}$, which can be used to represent stochastic mRNA synthesis and turnover [3], for instance. In this case, the protein synthesis rate $w_X^+ = V k_1^X r_X(t)$ depends on the number of mRNA molecules $r_X(t)$. The corresponding first- and second-order statistics of $r_X(t)$ at stationarity are given by

$$\langle r_X \rangle = \frac{\lambda_1}{\lambda_2} \tag{43}$$

$$\mathrm{Var}[r_X] = \frac{\lambda_1}{\lambda_2} = \sigma_X^2 \tag{44}$$

$$\kappa_X(\tau) = \sigma_X^2 e^{-\lambda_2 \tau}. \tag{45}$$

Ordinary differential equations capturing the time-evolution of these moments were derived using custom-made code in MATLAB (Version 9.10 (R2021a), The MathWorks Inc., Natick, Massachusetts) (`https://github.com/zechnerlab/bufferingvsnoise`). Probability densities over volume fractions were estimated by first sampling copy numbers ($N = 10^6$) from a multivariate normal distribution parameterizied by the LNA moments, and subsequently transformed into volume fractions.

#### S.1.5 Analytical results for the case of a two-component mixture

In the case of a two-component mixture ($\psi = 0$), we obtain approximate closed-form expressions that allow us to study concentration buffering and noise reduction analytically. This approximation considers small $\phi_1$ and $V_2$ and we refer to it as *dilute approximation.* At equilibrium, the two-component system can be described by the free energy

$$\begin{aligned} F(a_1, s_1) = v(a_1 n + s_1) f\left(\frac{a_1 n}{a_1 n + s_1}\right) + (V - v(a_1 n + s_1))\, f\left(\frac{(n - n_1) n v}{V - v(a_1 n + s_1)}\right) \\ + (36\pi)^{1/3} \gamma \left(V - v(a_1 n + s_1)\right)^{2/3}, \end{aligned} \tag{46}$$

with

$$f(\phi) = \frac{k_B T}{v}\Big[\chi_{AS}\phi(1-\phi) + \frac{\phi}{n}\log\phi + (1-\phi)\log(1-\phi)\Big]. \tag{47}$$

For simplicity, and because surface tensions tend to be low for biomolecular condensates [8], we focus on the limit of small surface tension and set $\gamma = 0$ in the following. The system exhibits two kinetic modes driven by the generalized thermodynamic forces that impose conditions on the rates

$$\log \frac{w_A^-(a_1, s_1)}{w_A^+(a_1 - 1, s_1 + n)} = -\frac{nv}{k_B T}(\mu_2^A - \mu_1^A) \tag{48}$$

$$\log \frac{w_S^-(a_1, s_1)}{w_S^+(a_1, s_1 - 1)} = -\frac{v}{k_B T}\left[\Pi_2 - \Pi_1\right], \tag{49}$$

where $\mu_\alpha^A = f'(\phi_\alpha)$ and $\Pi_\alpha = f(\phi_\alpha) - \phi_\alpha f'(\phi_\alpha)$. To simplify, we eliminate one of the two degrees of freedom by considering the system to be in osmotic equilibrium, such that $\Pi_1 = \Pi_2$ is satisfied at all times. Expressing $\Pi_1 = \Pi_2$ in terms of $\phi_1$ and $\phi_2$ yields

$$\frac{(\phi_1 - \phi_2)}{n}(n + n(\phi_1 + \phi_2)\chi_{AS} - 1) + \log(1 - \phi_1) - \log(1 - \phi_2) = 0. \tag{50}$$

This equation defines a function $\hat{\phi}_2 = f(\phi_1)$ that describes osmotic equilibrium. Unfortunately, this function cannot be expressed explicitly. We therefore focus on the dilute limit where $\phi_1$ is small. Considering only leading order terms, the free energy density in the dilute phase can be approximated as

$$f(\phi_1) \simeq \frac{k_B T}{v}\phi_1\left(\log\phi_1 - 1 + \chi_{AS}\right) \tag{51}$$

and (50) simplifies to

$$\frac{\phi_1}{n} + \frac{\phi_2}{n}(n + n\phi_2\chi_{AS} - 1) + \log(1 - \phi_2) \simeq 0. \tag{52}$$

For small $\phi_1$, however, the terms independent of $\phi_1$ will dominate such that $\hat{\phi}_2$ can be approximated by solving

$$\frac{\phi_2}{n}(n + n\phi_2\chi_{AS} - 1) + \log(1 - \phi_2) = 0 \tag{53}$$

with respect to $\phi_2$. As a result of this approximation, dense phase concentrations will be fixed to a value $\hat{\phi}_2$ that is independent of $\phi_1$. Eq. (53) is straightforward to solve numerically for a given set of parameters $n$ and $\chi_{AS}$. Once determined, we can express $s_1$ as a function of $a_1$ by solving

$$\hat{\phi}_2 = \frac{(a - a_1)nv}{V - v(a_1 n + s_1)}, \tag{54}$$

with respect to $s_1$, which yields

$$\hat{s}_1 = \frac{V}{v} - n\frac{a + a_1(\hat{\phi}_2 - 1)}{\hat{\phi}_2}. \tag{55}$$

Substituting this expression into the free energy (46) with $f(\phi_1)$ approximated by (51) and $\gamma = 0$ yields

$$F(a_1) = \frac{nv}{\hat{\phi}_2} f(\hat{\phi}_2)(a - a_1) + k_B T a_1 \left[ \log\left( \frac{a_1 v n}{V - nv(a - a_1)/\hat{\phi}_2} \right) - n \right] + k_B T \chi_{AS} a_1 n. \tag{56}$$

Defining $\xi = v f(\hat{\phi}_2)/(k_B T \hat{\phi}_2)$, we can write this more compactly as

$$\begin{aligned} \frac{F(a_1)}{k_B T} &= \xi(a - a_1)n + a_1 \left[ \log\left( \frac{a_1 nv}{V - nv(a - a_1)/\hat{\phi}_2} \right) - n \right] + \chi_{AS} a_1 n \\ &= -\mu(a - a_1)n + a_1 \left[ \log\left( \frac{a_1 nv}{V - nv(a - a_1)/\hat{\phi}_2} \right) - n \right] + const., \end{aligned} \tag{57}$$

with $\mu = \chi_{AS} - \xi$. Note that when $n = 1$ and $\hat{\phi}_2 = 1$, this reduced approximate system becomes equivalent to the binary system analyzed in our previous work [6]. This system can be further simplified in the case where the volume of the dense phase is small compared to the total volume ($V_2 \ll V$). In this case, the mixing entropy in (57) simplifies and we get

$$\frac{F(a_1)}{k_B T} \simeq -\mu(a - a_1)n + a_1 \left[ \log\left( \frac{a_1 nv}{V} \right) - n \right]. \tag{58}$$

Eq. (58) is the final form of the free energy that the dilute approximation is based on. Due to the simplifications made, only one kinetic mode remains, associated with the thermodynamic force

$$\begin{aligned} \log \frac{w_A^-(a_1)}{w_A^+(a_1 - 1)} &= \frac{F'(a_1)}{k_B T} \\ &= 1 - n + \mu + \log\left( \frac{a_1 nv}{V} \right). \end{aligned} \tag{59}$$

This force is zero when

$$a_{sat} = \frac{V}{nv} e^{-1+n-\mu} \tag{60}$$

or equivalently

$$\phi_{sat} = e^{-1+n-\mu}, \tag{61}$$

where we refer to $a_{sat}$ and $\phi_{sat}$ as *saturation concentration* and *saturation copy number*, respectively. This terminology reflects the fact that once the total volume fraction $\phi$ exceeds $\phi_{sat}$, the average dilute phase volume fraction $\langle \phi_1 \rangle$ remains constant at $\phi_{sat}$. The volume fraction $\phi_{sat}$ thus marks the threshold above which the mixture is saturated and phase coexistence is favorabe.

In the small droplet limit, we can use eq. (59) to express the partitioning rates as

$$w_A^-(a_1) = k_D a_1 \tag{62}$$

$$w_A^+(a_1 - 1) = w_A^-(a_1) e^{-\frac{F'(a_1)}{k_B T}} = k_D \frac{V}{nv} e^{1-n-\mu} = k_D a_{sat}, \tag{63}$$

where we have used $k_D = 6D/V^{2/3}$. In the considered regime, the exchange of solutes from the dilute phase to the dense phase is approximately linear, while the corresponding reverse rate becomes constant. In combination with solute synthesis and turnover, this system has degrees of freedom $(a_1, a)$, which evolve according to the events

$$(a_1, a) \underset{w_A^+}{\overset{w_A^-}{\rightleftharpoons}} (a_1 - 1, a) \tag{64}$$

$$(a_1, a) \underset{w_{A,1}^d}{\overset{w_{A,1}^b}{\rightleftharpoons}} (a_1 + 1, a + 1) \tag{65}$$

$$(a_1, a) \underset{w_{A,2}^d}{\overset{w_{A,2}^b}{\rightleftharpoons}} (a_1, a + 1). \tag{66}$$

Consistent with the small droplet limit, the synthesis rates in the dilute- and dense phase become $w^b_{A,1}(a_1, a) = Vk^A_1 r_A(t)$ and $w^b_{A,2}(a_1, a) = 0$, which means that solute synthesis in the dense phase becomes negligible. The turnover rates are given by $w^d_{A,1}(a_1, a) = k_2 a_1(t)$ and $w^d_{A,2}(a_1, a) = k_2(a(t) - a_1(t))$. Notice that because the rate $w^+_A$ is independent of $a(t)$, we can describe $a_1(t)$ in isolation by the effective birth-and-death process

$$a_1 \underset{Vk_1 r_A(t)+k_D a_{sat}}{\overset{(k_2+k_D)a_1}{\rightleftharpoons}} a_1 - 1, \tag{67}$$

or equivalently, $a_1(t) \sim \mathcal{BD}(Vk_1 r_A(t) + k_D a_{sat}, (k_2 + k_D)a_1(t))$. Here we have made use of the fact that two Poissonian event channels with the same stoichiometric change can be summarized into a single one with their respective rates added. Using an analogous derivation as for the generic birth-and-death process in Section S.1.3, we can write down the average solute copy number and noise strength at stationarity as

$$\langle a_1 \rangle = \frac{Vk_1\langle r_A\rangle + k_D a_{sat}}{k_2 + k_D} = \frac{k_D}{k_2 + k_D} a_{sat} + \frac{k_2}{k_2 + k_D}\langle a\rangle \tag{68}$$

$$\begin{aligned}\eta^2[a_1] &= \frac{k_2 + k_D}{Vk_1\langle r_A\rangle + k_D a_{sat}} + \left(\frac{Vk^A_1}{Vk_1\langle r_A\rangle + k_D a_{sat}}\right)^2 [k_2 + k_D]\,\hat{\kappa}(k_2 + k_D)\\ &= \frac{1}{\langle a_1\rangle} + \frac{1}{k_2 + k_D}\left(\frac{Vk^A_1}{\langle a_1\rangle}\right)^2 \hat{\kappa}(k_2 + k_D)\end{aligned} \tag{69}$$

Since $V_1 \to V$ in the limit of small droplets, the volume fraction $\phi_1$ is just $a_1$ times a constant, such that the statistics of $\langle \phi_1\rangle$ are readily obtained as

$$\langle \phi_1\rangle = \frac{k_D}{k_2 + k_D}\phi_{sat} + \frac{k_2}{k_2 + k_D}\langle \phi\rangle \tag{70}$$

$$\eta^2[\phi_1] = \frac{nv}{V}\frac{1}{\langle \phi_1\rangle} + \frac{1}{k_2 + k_D}\left(\frac{k^A_1 nv}{\langle \phi_1\rangle}\right)^2 \hat{\kappa}(k_2 + k_D), \tag{71}$$

where the second line is identical to (69) but expressed in terms of $\langle \phi_1\rangle$ instead of $\langle a_1\rangle$. Based on eq. (70), we can further define the apparent partition coefficient

$$\rho = \frac{\langle \phi_2\rangle}{\langle \phi_1\rangle} \simeq \frac{\hat{\phi}_2}{\langle \phi_1\rangle} = \frac{\hat{\phi}_2(1 + k_D/k_2)}{\phi_{sat}k_D/k_2 + \langle \phi\rangle}, \tag{72}$$

which decreases with $\langle \phi_1\rangle$ and $\langle \phi\rangle$.

#### S.1.6 Noise reduction in the limit of large volumes and quasi-static birth-rates

Based on the dilute approximation, noise reduction in the binary system is given by

$$\Gamma = \frac{\eta[\phi]}{\eta[\phi_1]} \simeq \sqrt{\frac{\frac{nv}{V}\frac{1}{\langle\phi\rangle} + \frac{1}{k_2}\left(\frac{k^A_1 nv}{\langle\phi\rangle}\right)^2 \hat{\kappa}(k_2)}{\frac{nv}{V}\frac{1}{\langle\phi_1\rangle} + \frac{1}{k_2+k_D}\left(\frac{k^A_1 nv}{\langle\phi_1\rangle}\right)^2 \hat{\kappa}(k_2 + k_D)}}. \tag{73}$$

To study how this relationship changes for macroscopic systems where intrinsic fluctuations of solute synthesis, turnover and partitioning become small, we take the large volume limit while keeping the diffusion rate $k_D$ constant. Fluctuations in the birth-rate $r_A(t)$ are considered to be independent of volume such that its stationary mean $\langle r_A\rangle$ and autocovariance function $\kappa(\tau)$ remain unaffected by volume scaling. If we now let $V \to \infty$, eq. (73) becomes

$$\begin{aligned}\lim_{V\to\infty}\Gamma &= \frac{\langle\phi_1\rangle}{\langle\phi\rangle}\sqrt{\frac{k_2 + k_D}{k_2}\frac{\hat{\kappa}(k_2)}{\hat{\kappa}(k_2 + k_D)}}\\ &= \underbrace{\left[1 + \frac{k_D}{k_2}\frac{\phi_{sat}}{\langle\phi\rangle}\right]}_{\simeq\Lambda}\sqrt{\frac{k_2}{k_2 + k_D}\frac{\hat{\kappa}(k_2)}{\hat{\kappa}(k_2 + k_D)}},\end{aligned} \tag{74}$$

where the first term in the second line is the buffering strength associated with the concentration dependency (70). Thus, noise reduction in the macroscopic limit can be approximated by the buffering strength multiplied by a factor, which depends on the autocovariance function $\kappa$ (or its Laplace transform $\hat{\kappa}$) and the parameters $k_2$ and $k_D$. Due to this factor, the noise reduction $\Gamma$ and the buffering strength $\Lambda$ can be substantially different, as reflected by our results in the main text. In the limit where fluctuations in $r_A(t)$ decay infinitely slowly, we have $\kappa(\tau) = \sigma^2$ and its Laplace transform becomes $\hat{\kappa}(u) = \sigma^2 u^{-1}$. Only in this special case, the second factor in (74) becomes equal to one such that noise reduction becomes equal to the buffering strength. This corresponds to the limit considered in ref. [9], where noise reduction has been studied based on equilibrium phase diagrams.

**Example.** To further illustrate the difference between concentration buffering and noise reduction, we analyze eq. (74) in the context of a simple example. In particular, we choose $\kappa(\tau) = \sigma_A^2 e^{-\lambda_2 \tau}$, corresponding to a stochastic process with a correlation time $\lambda_2^{-1}$. The Laplace transform of $\kappa$ is

$$\hat{\kappa}(u) = \frac{\sigma_A^2}{u + \lambda_2} \tag{75}$$

and correspondingly, (74) becomes

$$\Gamma = \left[1 + \frac{k_D}{k_2}\frac{\phi_{sat}}{\langle\phi\rangle}\right]\sqrt{\frac{k_2}{k_2 + k_D}\frac{k_2 + k_D + \lambda_2}{k_2 + \lambda_2}}. \tag{76}$$

As can be seen, noise reduction becomes identical to the buffering strength as $\lambda_2 \to 0$. For $\lambda_2 \to \infty$, we obtain the white noise limit, where

$$\Gamma = \left[1 + \frac{k_D}{k_2}\frac{\phi_{sat}}{\langle\phi\rangle}\right]\sqrt{\frac{k_2}{k_2 + k_D}}. \tag{77}$$

An interesting observation in eq. (77) is that for large timescale ratios $k_D/k_2$, noise reduction scales as $\sim \sqrt{k_D/k_2}$, whereas the buffering strength scales as $\sim k_D/k_2$. Thus, while both quantities approach infinity as $k_D/k_2$ becomes large, they do so at different rates.

### S.2 Analysis of previously published data

#### S.2.1 Synthetic system

We analyzed experimental concentration measurements from the 2NT-DDX4$^{\mathrm{YFP}}$ that we published previously using the theory outlined here [6]. In particular, we focussed on two datasets, one where dilute- and total concentrations $c_1$ and $c$ were measured for a large number of cells (Fig. 2c in ref. [6]), and one for which dilute- and dense phase concentrations $c_1$ and $c_2$ were measured for a smaller number of cells (Fig. S22 in ref. [6]). We used the former one to estimate timescale ratios using the calculated concentration dependency under the dilute approximation (70). To do so, we first estimated the relationship between $\langle c_1\rangle$ and $\langle c\rangle$ from the experimental measurements. To select cells well within the regime of phase coexistence, we excluded cells that had a total concentration below $12\mu M$. We divided the remaining measurements of total concentrations into $K$ non-overlapping intervals. The lower- and upper limit of the $k$th interval were determined as $l_k = q(k/(K+1))$ and $u_k = q((k+1)/(K+1))$ where $k = 0, \ldots, K-1$ and $q(x)$ is the empirical $x$-quantile of the experimentally measured sample of total concentrations. As an example, $q(0.5)$ would correspond to the median total concentration. The rationale behind using (equally-spaced) quantiles as the interval boundaries is that we obtain an approximately equal number of samples in each interval. Average total- and dilute phase concentrations for interval $k$, $\bar{c}^{(k)}$ and $\bar{c}_1^{(k)}$, were estimated using all cells $i$ for which $l_k < c^{(i)} \leq u_k$, where $c^{(i)}$ is the measured total concentration for cell $i$. The corresponding uncertainties of $\bar{c}^{(k)}$ and $\bar{c}_1^{(k)}$ were determined using bootstrapping [10].

Assuming that the number of samples in each interval is sufficiently large, we can make use of the central limit theorem and describe $(\bar{c}^{(k)}, \bar{c}_1^{(k)})^T$ as a Gaussian random vector such that

$$\begin{pmatrix}\bar{c}^{(k)}\\ \bar{c}_1^{(k)}\end{pmatrix} = \begin{pmatrix}\langle c\rangle\\ \langle c_1\rangle\end{pmatrix} + \begin{pmatrix}\Delta^{(k)}\\ \Delta_1^{(k)}\end{pmatrix} \quad \text{with} \quad \begin{pmatrix}\Delta^{(k)}\\ \Delta_1^{(k)}\end{pmatrix} \sim N(0, \Sigma^{(k)}) \quad \text{and} \quad \Sigma^{(k)} = \begin{pmatrix}\Sigma_{11}^{(k)} & \Sigma_{12}^{(k)}\\ \Sigma_{12}^{(k)} & \Sigma_{22}^{(k)}\end{pmatrix}, \tag{78}$$

where $\Sigma^{(k)}$ are the empirical uncertainties of $(\bar{c}^{(k)}, \bar{c}_1^{(k)})^T$ obtained via bootstrapping. Reformulating (70) in terms of concentrations (as opposed to volume fractions) and expressing $\langle c\rangle$ and $\langle c_1\rangle$ in terms of experimentally determined quantities $\bar{c}^{(k)}$ and $\bar{c}_1^{(k)}$ yields

$$\begin{aligned}\bar{c}_1^{(k)} &= \frac{k_D}{k_2 + k_D} c_{sat} + \frac{k_2}{k_2 + k_D}\bar{c}^{(k)} - \frac{k_2}{k_2 + k_D}\Delta^{(k)} + \Delta_1^{(k)}\\ &= \frac{k_D}{k_2 + k_D} c_{sat} + \frac{k_2}{k_2 + k_D}\bar{c}^{(k)} + \bar{\Delta}^{(k)},\end{aligned} \tag{79}$$

with $c_{sat} = \phi_{sat}/nv$ and $\bar{\Delta}^{(k)} = \Delta_1^{(k)} - k_2/(k_2 + k_D)\Delta^{(k)}$. Since $\bar{\Delta}^{(k)}$ is a linear combination of two Gaussians, it is itself a Gaussian, i.e., $\bar{\Delta}^{(k)} \sim N(0, \bar{\Sigma}^{(k)})$, with

$$\begin{aligned}\bar{\Sigma}^{(k)} &= \begin{pmatrix}-\frac{k_2}{k_2+k_D} & 1\end{pmatrix}\begin{pmatrix}\Sigma_{11}^{(k)} & \Sigma_{12}^{(k)}\\ \Sigma_{12}^{(k)} & \Sigma_{22}^{(k)}\end{pmatrix}\begin{pmatrix}-\frac{k_2}{k_2+k_D}\\ 1\end{pmatrix}\\ &= \frac{k_2^2}{(k_2 + k_D)^2}\Sigma_{11}^{(k)} + \Sigma_{22}^{(k)} - 2\frac{k_2}{k_2 + k_D}\Sigma_{12}^{(k)}.\end{aligned} \tag{80}$$

Thus, fitting the analytical concentration dependency to the experimentally determined one is a regression problem with parameter-dependent errors. Note that with $k_2$, $k_D$ and $c_{sat}$, the model is overparameterized and thus, not all parameters can be determined uniquely. However, we can reparameterize the model in terms of $k_D/k_2$ and $c_{sat}$ and thereby eliminate one unknown degree of freedom. The resulting two parameters and their uncertainties can be inferred using Bayes' rule, which reads

$$\begin{aligned}&p\left(k_D/k_2, c_{sat} \;\middle|\; (\bar{c}^{(0)}, \bar{c}_1^{(0)}), \ldots, (\bar{c}^{(K-1)}, \bar{c}_1^{(K-1)})\right)\\ &\propto p(k_D/k_2, c_{sat}) \prod_{k=0}^{K-1} N\left(\bar{c}_1^{(k)} - \frac{k_D}{k_2 + k_D} c_{sat} - \frac{k_2}{k_2 + k_D}\bar{c}^{(k)} \;\middle|\; 0, \bar{\Sigma}^{(k)}\right),\end{aligned} \tag{81}$$

where we consider the prior distribution $p(k_D/k_2, c_{sat})$ to be flat in the log-domain. Posterior distributions where sampled over logarithmic parameters (i.e., $\log k_D/k_2$ and $\log c_{sat}$) using the Metropolis-Hastings algorithm [11]. The proposal distribution was chosen to be a bivariate normal distribution such that $x^* \sim N(x, \sigma_p^2 \mathbb{I}_2)$ with $x$ and $x^*$ as the current and proposed parameters, respectively and $\mathbb{I}_2$ as the $2 \times 2$ identity matrix. The standard deviation $\sigma_p$ was chosen to be 0.1. The chain was simulated for $M = 10^5$ steps and the first $10^4$ samples were discarded to eliminate the initial burn-in period of the chain. Minimum mean squared error (MMSE) estimates were determined by calculating the component-wise mean of the resulting samples. Corresponding uncertainties were estimated as the component-wise standard deviations. When choosing the number of intervals to be $K = 34$, we estimated $c_{sat} = (7.471 \pm 0.053)\mu M$ and $k_D/k_2 = 21.569 \pm 1.426$. Doubling the number of intervals to $K = 68$ only marginally affected the inferred parameters ($c_{sat} = (7.494 \pm 0.050)\mu M$ and $k_D/k_2 = 22.560 \pm 1.4252$), showing that the results are robust for varying $K$.

We further analyzed partition coefficients in the 2NT-DDX4$^{YFP}$ system using the data from Fig. S22 in [6]. Rewriting eq. (72) in terms of concentrations yields for the apparent partition coefficient

$$\rho = \frac{\langle c_2\rangle}{\langle c_1\rangle} \simeq \frac{\hat{c}_2(1 + k_D/k_2)}{c_{sat}k_D/k_2 + \langle c\rangle}, \tag{82}$$

where $\langle c_2\rangle \simeq \hat{c}_2 = \hat{\phi}_2/(nv)$ under the dilute approximation. Thus, the partition coefficient is expected to decrease with average total (and thus, dilute phase-) concentration as long as $k_D/k_2$ is finite. As $k_D/k_2 \to$

$\infty$, the partition coefficient becomes $\hat{c}_2/c_{sat} = const.$ as the system approaches (quasi-)equilibrium. To compare these predictions to experiments, we calculated partition coefficients $c_2/c_1$ in individual cells. Since total concentrations were not quantified in this particular dataset, we plotted $c_2/c_1$ over $c_1$ as was done also in [12]. The resulting data is compared to the apparent partition coefficient $\rho = \hat{c}_2/\langle c_1 \rangle$, where $\hat{c}_2$ was determined as the average dense phase concentration calculated over individual cells (Supplementary Fig. S22).

#### S.2.2 Endogenous system

The data in Supplementary Fig. S.2b shows dilute- over total concentrations of fluorescently labelled NPM1 from two previously published studies [6, 12]. The first one corresponds to endogenously labelled NPM1 (Fig. 3f in ref. [6]. Total concentrations $c$ were measured during mitosis, when the nucleolus is dissolved. Dilute phase concentrations $c_1$ were measured ten hours later during interphase, when the nucleolus coexists with nucleoplasmic NPM1. Note that since $c$ and $c_1$ were measured at different time points, the resulting relationship between $c$ and $c_1$ is approximate. However, it nevertheless allows us to study concentration variability in the presence and absence of the nucleolus. To approximately convert fluorescence intensities to molar concentrations, we rescaled intensity values to achieve a mean total concentration of $7.7\mu M$ as measured previously [13]. The second dataset was obtained from [12], where fluorescently labelled NPM1 was overexpressed on top of native, unlabelled NPM1 (Fig. 1b in ref. [12]). These data were replotted as originally published and shown together with the calibrated, endogenous NPM1 measurements from our previous study [6] in Supplementary Fig. S.2b.

### S.3 Tables

Table S.1: Parameters used for the two-component system in Figure 2a and b

| Parameter | Value | Unit | Note |
|---|---|---|---|
| $V/v$ | $10^9$ | | |
| $n$ | 20 | | |
| $\gamma$ | $10^{-5}$ | $k_BT$ | |
| $\chi_{AS}$ | 1.2 | | |
| $V^{2/3}/(6D)$ | 154.72 | $s$ | Average diffusion time of $A$ |
| $k_1^A$ | * | $1/(Vs)$ | Varied to sweep over different $\langle \phi \rangle$ |
| $k_2$ | $2 \cdot 10^{-5}$ | $1/s$ | |
| $\lambda_1$ | $10^{-3}$ | $1/s$ | |
| $\lambda_2$ | $10^{-4}$ | $1/s$ | |
| $k_S$ | $5 \cdot 10^5$ | $1/s$ | |

Table S.2: Parameters used for the two-component system in Figure 2b inset

| Parameter | Value | Unit | Note |
|---|---|---|---|
| $V/v$ | $10^9$ | | |
| $n$ | 20 | | |
| $\gamma$ | $10^{-5}$ | $k_BT$ | |
| $\chi_{AS}$ | 1.2 | | |
| $V^{2/3}/(6D)$ | 154.72 | $s$ | Average diffusion time of $A$ |
| $k_1^A$ | $9.61 \cdot 10^{-8}$ | $1/(Vs)$ | |
| $k_2$ | $2 \cdot 10^{-5}$ | $1/s$ | |
| $\lambda_1$ | $10^{-3}$ | $1/s$ | |
| $\lambda_2$ | * | $1/s$ | Varied to sweep over different correlation times $1/\lambda_2$ |
| $k_S$ | $5 \cdot 10^5$ | $1/s$ | |

Table S.3: Parameters used for the three-component system in Fig. 2c and d

| Parameter | Value | Unit | Note |
|---|---|---|---|
| $V/v$ | $10^9$ | | |
| $n$ | 20 | | |
| $\gamma$ | $10^{-5}$ | $k_BT$ | |
| $\chi_{AS}$ | 0 | | |
| $\chi_{AB}$ | $-4$ | | |
| $\chi_{BS}$ | 0 | | |
| $V^{2/3}/(6D)$ | 92.83 | $s$ | Average diffusion time of $A$ and $B$ |
| $k_1^A$ | * | $1/(Vs)$ | Varied to sweep over different $\langle\phi\rangle$ |
| $k_1^B$ | $9 \cdot 10^{-8}$ | $1/(Vs)$ | |
| $k_2$ | $2 \cdot 10^{-5}$ | $1/s$ | |
| $\lambda_1$ | $10^{-3}$ | $1/s$ | |
| $\lambda_2$ | $10^{-4}$ | $1/s$ | |
| $k_S$ | $5 \cdot 10^5$ | $1/s$ | |

Table S.4: Parameters used for the two-component system in Supplementary Fig. S1

| Parameter | Value | Unit | Note |
|---|---|---|---|
| $V/v$ | $10^9$ | | |
| $n$ | 20 | | |
| $\gamma$ | $10^{-5}$ | $k_BT$ | |
| $\chi_{AS}$ | 1.2 | | |
| $V^{2/3}/(6D)$ | * | $s$ | Varied to achieve different timescale ratios $k_D/k_2$ |
| $k_1^A$ | * | $1/(Vs)$ | Varied to sweep over different $\langle\phi\rangle$ |
| $k_2$ | $5 \cdot 10^{-6}$ | $1/s$ | |
| $\lambda_1$ | $10^{-3}$ | $1/s$ | |
| $\lambda_1$ | $10^{-4}$ | $1/s$ | |
| $k_S$ | $5 \cdot 10^5$ | $1/s$ | |

Table S.5: Parameters used for the two-component system in Supplementary Fig. S3

| Parameter | Value | Unit | Note |
|---|---|---|---|
| $V/v$ | $10^9$ | | |
| $n$ | 20 | | |
| $\gamma$ | $10^{-5}$ | $k_BT$ | |
| $\chi_{AS}$ | 1.3 | | |
| $V^{2/3}/(6D)$ | 46.42 | $s$ | Average diffusion time of $A$ |
| $k_1^A$ | * | $1/(Vs)$ | Varied to sweep over different $\langle\phi\rangle$ |
| $k_2$ | $5 \cdot 10^{-6}$ | $1/s$ | |
| $\lambda_1$ | $3 \cdot 10^{-3}$ | $1/s$ | |
| $\lambda_2$ | $5 \cdot 10^{-4}$ | $1/s$ | |
| $k_S$ | $5 \cdot 10^5$ | $1/s$ | |

### References

[1] Yuichi Taniguchi, Paul J Choi, Gene-Wei Li, Huiyi Chen, Mohan Babu, Jeremy Hearn, Andrew Emili, and X Sunney Xie. Quantifying e. coli proteome and transcriptome with single-molecule sensitivity in single cells. *Science*, 329(5991):533–538, 2010.

[2] Arren Bar-Even, Johan Paulsson, Narendra Maheshri, Miri Carmi, Erin O'Shea, Yitzhak Pilpel, and Naama Barkai. Noise in protein expression scales with natural protein abundance. *Nature*

*genetics*, 38(6):636–643, 2006.

[3] Johan Paulsson. Summing up the noise in gene networks. *Nature*, 427(6973):415–418, 2004.

[4] Brian Munsky, Gregor Neuert, and Alexander Van Oudenaarden. Using gene expression noise to understand gene regulation. *Science*, 336(6078):183–187, 2012.

[5] Joel L Schiff. *The Laplace transform: theory and applications*. Springer Science & Business Media, 2013.

[6] Adam Klosin, Florian Oltsch, Tyler Harmon, Alf Honigmann, Frank Jülicher, Anthony A Hyman, and Christoph Zechner. Phase separation provides a mechanism to reduce noise in cells. *Science*, 367(6476):464–468, 2020.

[7] Nicolaas Godfried Van Kampen. *Stochastic processes in physics and chemistry*, volume 1. Elsevier, 1992.

[8] Clifford P Brangwynne, Christian R Eckmann, David S Courson, Agata Rybarska, Carsten Hoege, Jöbin Gharakhani, Frank Jülicher, and Anthony A Hyman. Germline p granules are liquid droplets that localize by controlled dissolution/condensation. *Science*, 324(5935):1729–1732, 2009.

[9] Dan Deviri and Samuel A Safran. Physical theory of biological noise buffering by multicomponent phase separation. *Proceedings of the National Academy of Sciences*, 118(25):e2100099118, 2021.

[10] B. Efron. Bootstrap Methods: Another Look at the Jackknife. *The Annals of Statistics*, 7(1):1 – 26, 1979.

[11] Christian P Robert, George Casella, and George Casella. *Monte Carlo statistical methods*, volume 2. Springer, 1999.

[12] Joshua A Riback, Lian Zhu, Mylene C Ferrolino, Michele Tolbert, Diana M Mitrea, David W Sanders, Ming-Tzo Wei, Richard W Kriwacki, and Clifford P Brangwynne. Composition-dependent thermodynamics of intracellular phase separation. *Nature*, 581(7807):209–214, 2020.

[13] Brian D Freibaum, James Messing, Peiguo Yang, Hong Joo Kim, and J Paul Taylor. High-fidelity reconstitution of stress granules and nucleoli in mammalian cellular lysate. *Journal of Cell Biology*, 220(3):e202009079, 2021.
